# NeuroMabSeq: high volume acquisition, processing, and curation of hybridoma sequences and their use in generating recombinant monoclonal antibodies and scFvs for neuroscience research

**DOI:** 10.1101/2023.06.28.546392

**Authors:** Keith G. Mitchell, Belvin Gong, Samuel S. Hunter, Diana Burkart-Waco, Clara E. Gavira-O’Neill, Kayla M. Templeton, Madeline E. Goethel, Malgorzata Bzymek, Leah M. MacNiven, Karl D. Murray, Matthew L. Settles, Lutz Froenicke, James S. Trimmer

## Abstract

The Neuroscience Monoclonal Antibody Sequencing Initiative (NeuroMabSeq) is a concerted effort to determine and make publicly available hybridoma-derived sequences of monoclonal antibodies (mAbs) valuable to neuroscience research. Over 30 years of research and development efforts including those at the UC Davis/NIH NeuroMab Facility have resulted in the generation of a large collection of mouse mAbs validated for neuroscience research. To enhance dissemination and increase the utility of this valuable resource, we applied a high-throughput DNA sequencing approach to determine immunoglobulin heavy and light chain variable domain sequences from source hybridoma cells. The resultant set of sequences was made publicly available as searchable DNA sequence database (<u>neuromabseq.ucdavis.edu</u>) for sharing, analysis and use in downstream applications. We enhanced the utility, transparency, and reproducibility of the existing mAb collection by using these sequences to develop recombinant mAbs. This enabled their subsequent engineering into alternate forms with distinct utility, including alternate modes of detection in multiplexed labeling, and as miniaturized single chain variable fragments or scFvs. The NeuroMabSeq website and database and the corresponding recombinant antibody collection together serve as a public DNA sequence repository of mouse mAb heavy and light chain variable domain sequences and as an open resource for enhancing dissemination and utility of this valuable collection of validated mAbs.

## Introduction

Using antibodies (Abs) to detect endogenous target proteins in brain samples is foundational to many aspects of neuroscience research. Antibodies provide specific and effective labeling of endogenous targets in diverse brain samples including those obtained from human donors ^1^. Antibody labeling can be detected with various imaging modalities, allowing for determination of spatial details of protein expression and localization across a wide range of scales, which in neuroscience research can range from single molecules to nanoscale molecular assemblies to cells to intact brain circuits ^1^. Conventional (i.e., non-recombinant) Abs can be produced in a variety of animal species (e.g., mice, rats, rabbits, goats, etc.) as polyclonal Abs, and from hybridoma cell lines as monoclonal Abs (mAbs) ^2^. Combinatorial (i.e., multiplex) labeling and detection can be performed using combinations of Abs from different species followed by their detection with species-specific dye-conjugated secondary Abs. Moreover, except for those of rabbit origin, within a given species individual mAbs exist as one of multiple IgG subclasses, and those of different IgG subclasses can be multiplexed and separately detected with subclass-specific secondary Abs ^3^.

As a consequence of mAb development efforts that span over 30 years, including at the UC Davis/NIH NeuroMab Facility, we have generated a large collection of cryopreserved hybridoma cells producing mouse mAbs. These mAbs have well-defined target specificities and efficacies for immunolabeling endogenous target proteins in mammalian brain samples by immunoblot (IB) and immunohistochemistry (IHC) applications ^4–6^. Cryopreserved archives of viable mAb-producing hybridoma cells define mAbs as renewable research reagents, a major distinguishing characteristic of mAbs when compared to polyclonal Abs ^7^. However, the continued availability of a given mAb is not absolutely guaranteed as it relies on the successful recovery into cell culture of these cryopreserved hybridoma cells, and that these cells in culture continue to reliably produce the exact same mAb that was characterized during its development.

The target binding specificity and efficacy of a given Ab is defined by its light and heavy chain variable domains (i.e., V_L_ and V_H_ domains) that together with the light and heavy chain constant regions define the full Ab molecule ^2^. Determining the sequence of a particular mAb’s V_L_ and V_H_ domain generates a truly permanent and unique Ab archive in the form of DNA sequence ^8^. Furthermore, utilizing such sequence information to generate plasmids expressing recombinant forms of these mAbs (R-mAbs) effectively eliminates the need for the expensive and labor-intensive maintenance of cryopreserved hybridoma collections in liquid nitrogen and allows for inexpensive archiving and simple dissemination as nucleotide sequence and/or plasmid DNA. Defining the primary sequence of mAbs also allows for their use as molecularly defined research regents, enhancing their value in terms of research transparency ^8^. Recombinant expression can also afford more reliable and often higher-level expression than from hybridomas and enhance research reproducibility as the expression plasmid can be resequenced prior to each use ^8^. Plasmids can also be archived at and disseminated from open access nonprofit resources such as Addgene (https://www.addgene.org/), with increased ease and lower cost dissemination than cryopreserved hybridoma cells. Cloning and recombinant expression also allows for diverse forms of Ab engineering. This includes engineering to confer distinct detection modalities to the expressed mAb, facilitating their use in multiplex labeling ^9^, as well as development of miniaturized Abs such as single chain variable region fragments (scFvs) ^10, 11^ with additional advantages due to their small size, which enhances tissue penetration and allows for increased imaging resolution ^12^.

To generate a lasting archive and obtain recombinant Abs with enhanced opportunities for engineering, we sequenced the V_L_ and V_H_ domains of mAbs in our large and extensively characterized collection. Initial efforts used RT-PCR-based cloning of mAb V_L_ and V_H_ domains into mammalian expression plasmids followed by Sanger plasmid sequencing. This led to the successful cloning, sequencing, and expression of almost 200 of our mAbs ^9^, but this effort only represented a small fraction of the ≈2,400 mAbs in our extensive collection. Here we describe the development of a pipeline for high-throughput sequencing of hybridomas to obtain mAb V_L_ and V_H_ domain sequences. We also detail novel bioinformatics approaches used to analyze the quality of the obtained sequences and the diversity of identified V_L_ and V_H_ domain sequences. Together these efforts have led to a large public repository of V_L_ and V_H_ domain sequences. We also used these sequences to generate R-mAb expression plasmids that are available through open access resources. We also describe pipelines for engineering these R-mAbs into forms with distinct detection modalities and miniaturizing them into scFvs. Together these efforts have generated a resource that further enables antibody-based neuroscience research and serve as a model for enhancing the archiving and dissemination of other mAb collections in recombinant form.

## Results

Our mAb development projects typically start with 960-2,880 candidate oligoclonal hybridoma samples, from a set of between 10-30 x 96 well microtiter plates in which the initial products of the mouse splenocyte-myeloma fusion reaction are cultured ^5^. These cultures and the Abs they produce are oligoclonal, likely containing more than one hybridoma clone, but producing a collection of Abs much less complex than that present in polyclonal antiserum and/or affinity-purified polyclonal Ab preparations. We refer to these hybridoma samples as “parent” samples as it is from these initial oligoclonal samples that monoclonal hybridomas and mAbs are derived by subcloning to monoclonality. Conditioned medium from each culture well, referred to as tissue culture supernatants or TC supes, is evaluated by ELISA from which we typically identify 24-144 ELISA positive hybridoma samples for expansion and further characterization. The TC supes from each of these expanded parent hybridoma cultures are subsequently evaluated by numerous assays (transfected cell immunocytochemistry/ICC, brain immunohistochemistry/IHC, and brain immunoblots/IB being the standard set) in parallel ^4–6^. A subset of parent hybridomas, up to five per project, are selected for subcloning to monoclonality by limiting dilution ^2^. We typically retain and archive five independent target-positive subclones for each parental hybridoma cell line with the expectation that these are independent isolates of a single clone of target-positive hybridoma cells present in the oligoclonal parent hybridoma culture. Relatively few target-positive wells (e.g., 5%) are observed among the large collection of parent samples initially screened ^2, 5^, suggesting that it is unlikely that there exist more than one target-positive hybridoma clone in the oligoclonal parental cell culture.

Our mAb nomenclature reflects this process. In our naming system, individual mAb projects are designated by a letter (for the most part a K, L or N) and a number, followed by a “/” and the number assigned to that ELISA-positive sample. For example, for project K89, performed in 2001 and targeting the Kv2.1 voltage-dependent K^+^ channel, we cryopreserved 48 parent hybridomas samples, designated K89/1-K89/48. We selected three of these parental lines for subcloning, one of them being K89/34. The five subclones derived from the K89/34 parent were designated K89/34.1-K89/34.5, for which multiple vials of each were cryopreserved. These monoclonal K89/34 cell lines have been expanded numerous times in the past 20 years, such that in total our cryopreserved hybridoma collection contains over 50 vials of the various K89/34 subclones archived at different times.

Overall, from ≈800 mAb projects, we have ≈45,000 distinct parent oligoclonal hybridoma cell lines, comprising a collection of ≈60,000 cryopreserved vials of parent oligoclonal hybridomas. From these parents, we subcloned by limiting dilution ≈3,500 distinct parental hybridoma cell lines, from which we cryopreserved between 1-5 subcloned monoclonal sample per parent, for a total of ≈11,000 distinct samples. As we typically cryopreserve multiple vials of individual subclones, our collection comprised ≈26,000 vials of cryopreserved monoclonal hybridoma samples. For sequencing purposes, which focused on monoclonal samples, we separated these samples into two classes. The first we defined as “biological replicates”, representing independent subclones obtained from the subcloning of a single parental hybridoma culture (e.g., K89/34.1 and K89/34.2 are biological replicates of the K89/34 hybridoma). The second class we defined as “technical replicates”, comprising independent cryopreserved vials of the same subcloned hybridoma (e.g., distinct vials of K89/34.1, including those cryopreserved on different dates). Our overarching goal was to use the novel high throughput sequencing and bioinformatics workflow we developed (Figure 1), as detailed below, to obtain corroborating V_L_ and V_H_ domain sequences from at least three biological replicates for each of the ≈3,500 subcloned mAbs in our collection.

**Figure 1.**
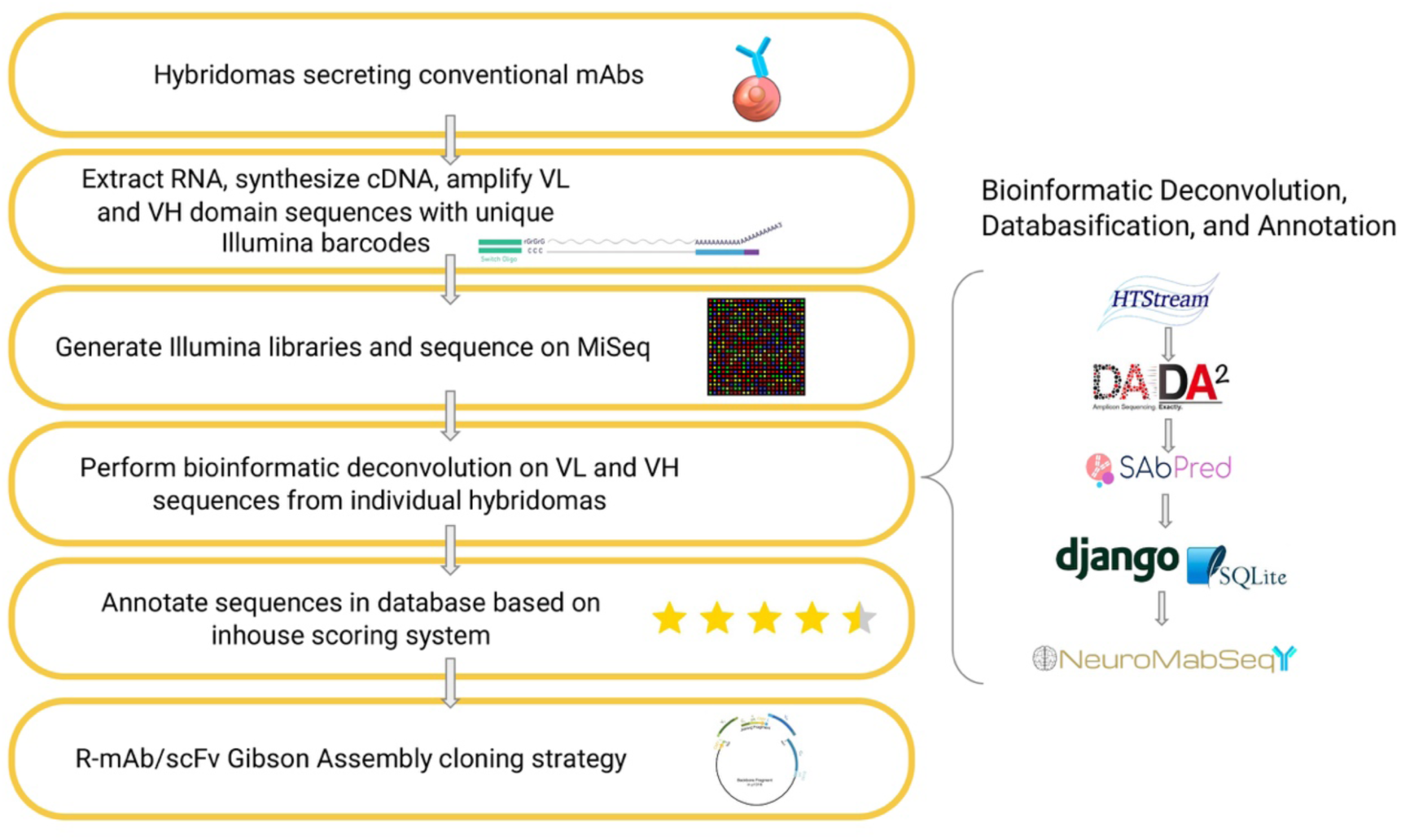
Sequencing workflow and bioinformatics processing. Hybridomas of interest are sequenced using a workflow consisting of RNA extraction, cDNA synthesis, and semi-nested PCR amplification with IgG-specific primers followed by the addition of unique Illumina barcodes to each sample. Illumina libraries are then generated and adapters are ligated for sequencing on the MiSeq platform. Bioinformatics processing is shown on the right panel. Reads from the Illumina sequencing are run through HTStream for base quality trimming and other read processing. Next, they are passed through DADA2 for amplicon denoising followed by SAbPred ANARCI tool based on the IMGT numbering scheme. All ASVs, metadata, and other quality metrics are uploaded to the NeuroMabSeq database and website where further information and tools are provided to the end users. This includes but is not limited to BlastIR results, BLAT searches across the database, and recommended high quality sequences for recombinant antibody design. Annotations of internally generated scores are provided in addition to other database information. Finally, high quality sequences are used in the design of gene fragments for generation of R-mAb and scFv expression plasmids.

### Establishment of a hybridoma sequencing pipeline

Prior to initiating large-scale sequencing efforts, we optimized the sequencing pipeline, beginning with processing of the frozen collection of hybridoma cells, and all subsequent steps, up to and including Illumina MiSeq sequencing (Figure 1). We previously found that RNA of sufficient quantity and quality for RT-PCR based cloning of V_L_ and V_H_ domain sequences could be isolated directly from cryopreserved hybridoma cells, without the need to recover the cells into culture ^9^. As such, we assumed that this would also hold for obtaining RNA that would enable effective and reliable sequencing of the mAb V_L_ and V_H_ domains employing Illumina-based high throughput sequencing. We made aliquots of hybridoma cells in 96 well plates after rapid thawing and after a single PBS wash, lysed them and isolated RNA using a QiaCube HT system. RNA was quantified on a well-by-well basis by Nanodrop readings and normalized across all wells of the plate to a range of 7-15 ng/µL.

### Sequencing of antibody variable regions

The sequencing library preparation employed a 5’-RACE like approach combined with a semi-nested barcode-indexing PCR (Supplementary Figure 1). The protocol of Meyer, DuBois, and colleagues ^13^ was modified to reverse transcribe four transcripts in a single reaction, employing a cocktail of four reverse transcription primers (see Supplementary Table 1 for all primer sequences). Two of these reverse primers were specific for the mouse heavy chain constant region, one representing a sequence conserved in the heavy chain constant regions of the IgG1, IgG2a and IgG2b subclasses, and the other specific for the IgG3 subclass. The second pair of reverse primers used were specific for the mouse kappa and lambda light chain constant region, respectively. We also utilized a shorter version of the template switching oligo (TSO) than used previously ^13^ to preserve more sequencing cycles for the regions of interest. The cDNA was subsequently PCR-amplified with a cocktail of four nested constant region chain-specific reverse primers analogous but internal to those used in the cDNA synthesis reaction on the 3’ end, and barcode-indexed forward primers, targeting the TSO sequence, on the 5’ end (Supplementary Figure 1). This resulted in incorporation of 96 unique inline barcode indices that were used to uniquely identify each well of a source sample plate. To create sequence diversity for the subsequent Illumina sequencing step, the lengths of the inline indices varied between five to eight bases, staggering the readthrough through the TSO sequence shared by all amplicons. A representative subset of the PCR products was checked for quality via microcapillary electrophoresis. After pooling, the amplicons were converted into sequencing libraries by ligation of Illumina adapters. Sets of barcoded amplicons from each 96 well plate were pooled and cleaned up with SPRI-beads. Each pool was subsequently converted into one Illumina-barcode indexed sequencing library using the ThruPLEX DNA-Seq HV kit from Takara Bio. Libraries of up to twelve 96-well plates were sequenced on one MiSeq run with paired-end 300 bp sequencing read to provide overlap for the region of interest.

### Bioinformatics processing

A novel bioinformatic pipeline was developed to analyze the resultant sequences and make them easily and publicly accessible via a software package, database, and website (Figure 1). The forward and reverse reads from the Illumina sequencing were joined bioinformatically, and demultiplexed to the sample level using Illumina barcodes and Illumina bcl2fastq (v. 2.20) software. Primer sequence was used to determine whether the sequence obtained corresponded to mouse V_L_ or V_H_ and was then removed. TSO sequence was identified and removed, any sequence containing a ‘N’ character was removed from further consideration, low quality base pairs (<10 q-values) were removed from the 3’ ends, followed by overlapping of paired reads using HTStream (https://github.com/ibest/HTStream, v. 1.2.0-release). Overlapping reads that met a minimum length threshold of 385 bp were then denoised and summarized into Amplicon Sequence Variants (ASVs) using the DADA2 algorithm ^14^ and filtered using a custom R script (Figure 1).

We next applied a filter to remove sequences corresponding to the aberrant/non-productive V_L_ transcript present in Sp2/0 hybridomas. All hybridomas in our collection were developed from the fusion of Balb/c mouse splenocytes with the Sp2/0-Ag14 (Sp2/0; ATCC CRL-1581) myeloma cell line ^4, 5^. The Sp2/0 myeloma cell line has numerous advantages for hybridoma development ^15^. However, Sp2/0 cells are themselves hybridomas, derived from the MOPC-21 myeloma cell line ^14^. MOPC-21 and therefore Sp2/0 cells express a non-productive (i.e., untranslated) mutated aberrant kappa IgG light chain transcript ^16^. We eliminated ASVs containing these sequences so that they were not included in any subsequent analyses.

We then applied a filter to eliminate sequences that did not correspond to the full-length coding regions of mouse V_L_ and V_H_ domains. The nucleotide sequences were first translated *in silico*, and the deduced amino acid sequences were analyzed for correspondence to expected immunoglobulin sequences using the ANARCI (Antigen receptor Numbering And Receptor ClassificatIon) tool as applied to Ab variable domains ^17^ downloaded from the SAbPred structure-based Ab prediction server (https://opig.stats.ox.ac.uk/webapps/sabdab-sabpred/sabpred/) ^18^. This tool defined the sequences corresponding to the expected primary structure of the V_L_ and V_H_ domains. It assigned each amino acid a position corresponding to the IMGT (international ImMunoGeneTics information system) numbering scheme ^19^. This entailed aligning the obtained sequence against that of consensus V_L_ and V_H_ domains and assigning each amino acid a slot within IMGT positions a.a. 1-127 of the light chain and 1-128 of the heavy chain, inserting gaps or extra positions (e.g., 112A, 112B, etc.) when needed to maintain the alignment of the obtained sequence to the IMGT consensus positions. The full translated sequence and the amino acid sequence corresponding to that within the boundaries of the V_L_ and V_H_ domains are displayed on our publicly available NeuroMabSeq website (https://neuromabseq.ucdavis.edu/) (Figure 2).

**Figure 2.**
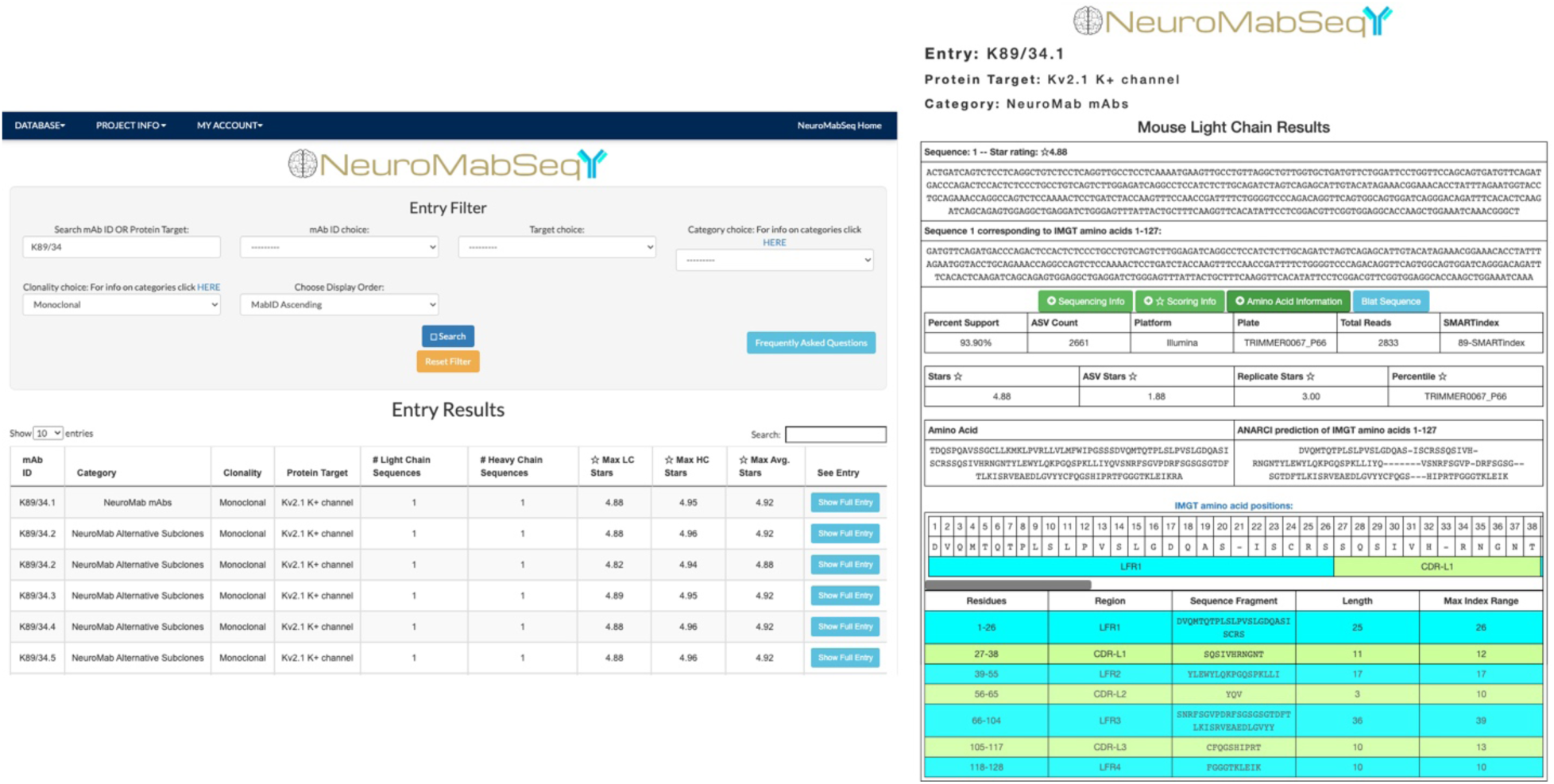
Example of a NeuroMabSeq website sequence entry. (a) View of the query interface that the website provides where users can search by mAb ID, target, etc. Shown are sequences returned for a search for “K89/34”. (b) The entry for the K89/34.1 light chain is shown with separate boxes for “Sequencing Information ’’, “Scoring Information” and “Amino Acid Information”. The “Sequencing Information’’ dropdown contains data such as the number of ASVs attributed to the obtained sequence and the number of total reads attributed to light chains or heavy chains for the sample, as well as the plate. The “Scoring Information” reveals the star rating assigned to each sequence, as well as the contribution of the ASV-based star and replicate-based star components of the scoring to the total score. The “Amino Acid Information” dropdown contains information such as the full amino acid sequence, the sequence corresponding to the ANARCI prediction of IMGT amino acid positions for the V_L_ domain, and within this the FR 1-4 and CDR 1-3 boundaries. The nucleotide sequence corresponding to the ANARCI prediction of IMGT amino acids is also shown to facilitate design of gBlocks for Gibson Assembly-based cloning of recombinant mAbs and scFvs. In addition, the “BLAT Sequence” feature is available to compare this sequence to all other sequences in the database. An analogous set of information is supplied for the heavy chain.

Next, translated V_L_ and V_H_ domain sequences were aligned to framework regions (FR) and complementarity determining regions (CDR) using the Abysis tool http://www.abysis.org/. This allowed for the generation of trimmed amino acid sequences corresponding to only the V_L_ and V_H_ domains themselves. These sequences were assigned positions with the FR and CDR domains using the Abysis tool, which is displayed in a color-coded format on the NeuroMabSeq website (Figure 2). Translated hybridoma sequences that did not yield any amino acids within any one of the assigned FR1-4 or CDR1-3 regions were filtered from the database, as were any sequences lacking the first 10 amino acids of FR1 or the last 10 amino acids of FR4. The original nucleotide sequence was then trimmed to remove sequences outside of that encoding IMGT positions a.a. 1-127 of the light chain and 1-128 of the heavy chain and shown in a separate box on the NeuroMabSeq website (Figure 2), facilitating their subsequent use in the development of R-mAb and scFv expression plasmids. *Bona fide* V_L_ and V_H_ encoding nucleotide sequences remaining after these filtering steps were quantified using amplicon sequence variant (ASV) analysis ^14, 20^.

Finally, sequences from each hybridoma sample were filtered based on ASV count and only those sequences corresponding to ≥10% of the total ASVs were included in the database. The ASV counts, quality scoring, and amino acid information of each distinct sequence corresponding to IMGT amino acids are grouped additively and displayed on the NeuroMabSeq website (Figure 2). This allows for ease of access to all nucleotide information, amino acid information, and tools embedded on the website.

To date we have sequenced 8,642 novel (i.e., non-control) monoclonal hybridoma samples using this approach (Figure 3). Of these, 1,903 samples (22%) did not yield any sequences that met the sequencing criteria, including any sequences for the aberrant Sp2/0-derived V_L_ domain (referred to as “dropout” samples). Removing these yielded a database of 6,739 hybridoma samples, which yielded 15,064 distinct ASVs. After eliminating those that did not correspond to bona fide V_H_ and V_L_ sequences, 13,401 were used as the input to the star scoring system (see details below). Finally, sequences were grouped by whether they came from biological or technical replicates of hybridomas with the same mAb ID (Figure 3).

**Figure 3.**
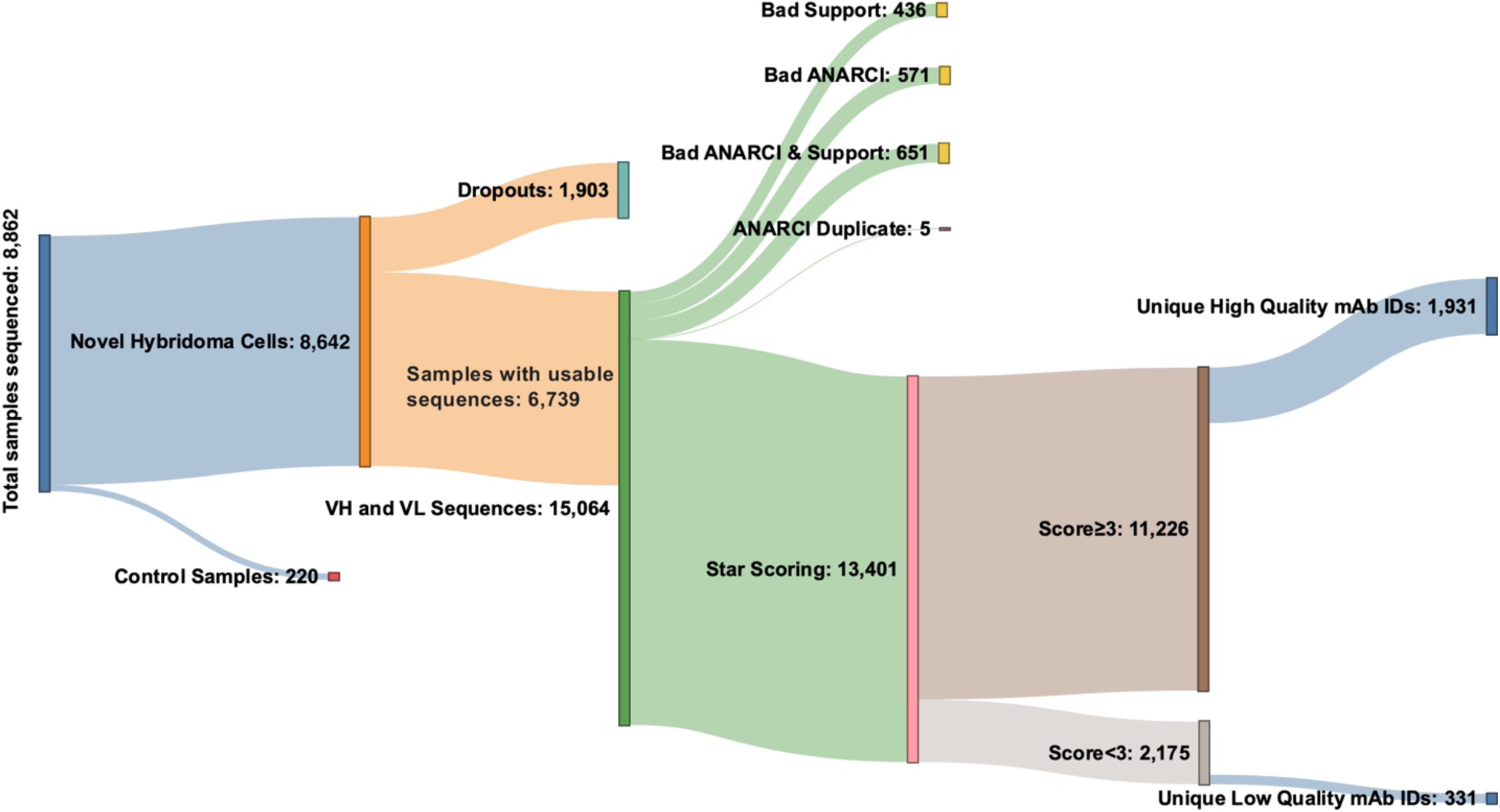
NeuroMabSeq workflow with details of samples sequenced. This Sankey diagram depicts the details of hybridoma samples sequenced to date. The 8,642 novel (i.e., non-control hybridoma) samples sequenced included 1,903 that did not yield usable sequences and were designated as “dropouts”. 6,739 samples returned usable sequences that yielded 15,064 total V_L_ and V_H_ sequences. 13,401 of these remained after eliminating any sequences that had insufficient support, did not conform to ANARCI conventions of valid antibody sequences, or were duplicates; these were subjected to the star scoring system. Of these, 11,226 had sequencing quality scores greater than 3, while 2,175 samples had sequencing quality scores less than 3. The number of unique mAb IDs after grouping all biological and technical replicates is also provided (1,931 high quality and 331 low quality).

### Representation of monoclonal hybridomas that express multiple productive V_L_ and V_H_ domain transcripts

Prior studies have revealed the presence of more than one productive light and heavy chain transcript in a subset of what are presumably monoclonal mouse and rat hybridoma cells ^21–27^. The largest study to date used sequence information, primarily from Sanger sequencing of plasmids containing V_L_ and V_H_ domains obtained from PCR-based cloning, to evaluate the incidence of additional productive V_L_ and V_H_ transcripts in 185 otherwise unrelated rat and mouse hybridoma cell lines from seven different laboratories ^27^, many of which produce mAbs that are commercially available. Approximately one-third of these hybridomas (59/185) were found to contain an additional productive (i.e., non-aberrant) light and/or heavy chain transcript^27^. Our large-scale sequencing efforts represent a valuable opportunity to determine the prevalence of V_L_ and V_H_ domain sequences in a much larger set of curated sequencing samples from 1,931 distinct monoclonal mouse hybridomas. We found that in addition to the predominant sequence with the preponderance of ASV counts, a substantial subset of our samples contain low levels of one or more alternate productive V_L_ and/or V_H_ domain sequences that represent <10% of the total ASV count (Figure 4). It is not clear due to their relatively low abundance whether these sequences represent biologically relevant transcripts (i.e., would substantively impact the population of mAb produced by that hybridoma). However, another subset of samples contains alternate sequences, primarily VL, at levels similar to the predominant component of the total ASV counts returned (e.g., representing 30-50% of the ASV counts), and presumably would make a substantive contribution to the Abs produced from that hybridoma. From the 1,931 unique high quality monoclonal hybridoma cell lines we sequenced that contained a bona fide V_H_ and VL, we found that 37 (1.9%) consistently contained, across biological and technical replicates, an additional productive (i.e., non-aberrant) V_L_ and/or V_H_ transcript that represented ≥10% of the total count of productive V_L_ or V_H_ encoding reads corresponding to the ASVs (Table 1). We analyzed whether the representation of these values differed based on whether the hybridomas were made during the earlier period when PEG chemical fusion was used to generate hybridomas, or the more recent period when electrofusion was used ^5^. We did not detect any difference in the overall representation of V_H_ or V_L_ transcripts between unique mAb sequences from these two periods (Table 1).

**Figure 4.**
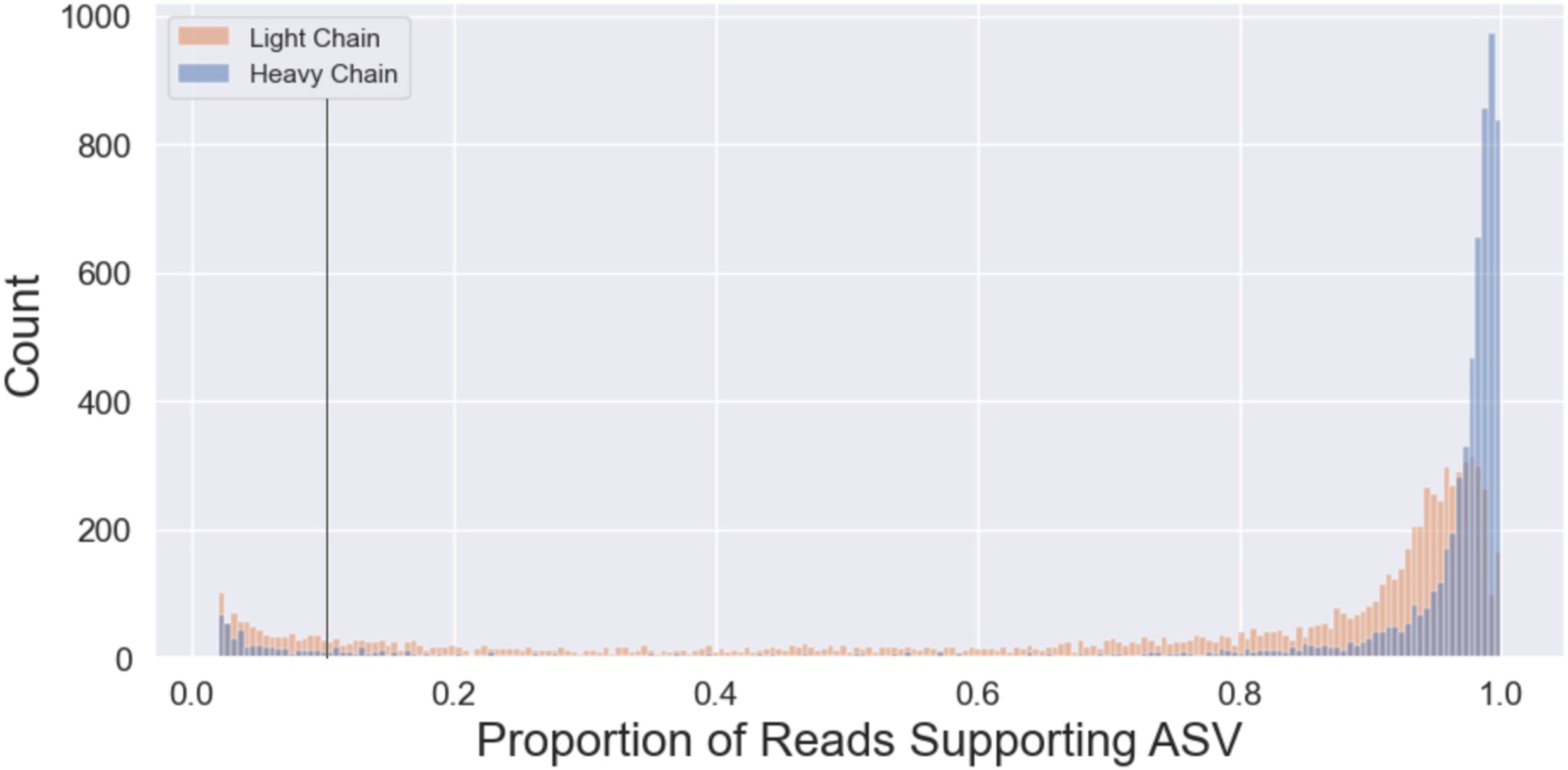
ASV Percent Score reported for the V_L_ and V_H_ sequencing for each of the samples sequenced. Based on these distributions, and assumptions as to unlikely biological contributions of very minor transcripts, a cut off was applied so that only ASVs representing >10% of total reads (black line) were considered.

**Table 1.** Sequencing summary after scoring and filtering.

| Category | Total Count | Total Percent | Chemical Fusion Count | Chemical Fusion Percent | Electrofusion Count | Electrofusion Percent |
| --- | --- | --- | --- | --- | --- | --- |
| $V_L$ or $V_H = 0$ | 341 | 17.7% | 186 | 17.2% | 155 | 18.3% |
| Total ($V_L \geq 1$ and $V_H \geq 1$) | 1590 | 82.3% | 896 | 82.8% | 694 | 81.7% |
| No additional chain | 1553 | 80.4% | 874 | 80.0% | 679 | 80.0% |
| Additional chains of some sort | 37 | 1.9% | 22 | 2.0% | 15 | 1.8% |
| One additional chain of some sort | 37 | 1.9% | 22 | 2.0% | 15 | 1.8% |
| One additional $V_L$ only | 23 | 1.2% | 14 | 1.3% | 9 | 1.1% |
| One additional $V_H$ only | 4 | 0.2% | 2 | 0.2% | 2 | 0.2% |
| One additional $V_L$ and $V_H$ | 10 | 0.5% | 6 | 0.6% | 4 | 0.5% |

**Table 1**. Sequencing summary after scoring and filtering. Total number of high quality unique mAb ID sequences (1,931 total) from projects performed using PEG chemical fusion (1,082 total) and electrofusion (849 total) to generate the hybridomas. Samples with at least one V_L_ and one V_H_ sequence are then separated into those containing exactly one V_L_ and one V_H_ sequence, one additional chain of some sort, one additional VH, one additional VL, and one additional V_L_ and VH. Projects performed using either PEG chemical fusion or electrofusion were then analyzed separately.

### Development of a scoring system to rank the quality of sequences from individual hybridoma samples

To rank the quality of sequence support for any given mAb and identify the best sequence to utilize in subsequent cloning efforts, we developed a simple scoring system. Scoring was based on both the quantity of sequences as defined by ASV counts, and the quality of the sequences obtained. This is defined by the presence of that sequence among biological and technical replicate samples of the same hybridoma, and the lack of that sequence in unrelated hybridoma samples (see Methods for details). The scoring system returned scores on a continuous scale ranging from 0-5 based on read support, defined as the ratio of a particular ASV relative to total reads produced for that sample, as well as consistency across biological and technical replicates of the same hybridoma cell line. The result of this scoring system and the distribution of V_L_ and V_H_ scores is shown in Figure 5. Individual V_L_ sequences often have a lower proportion of total ASV read support compared to V_H_ sequences (Figure 5), likely due to the penalty imposed because of the higher incidence of samples yielding multiple V_L_ sequences than seen for V_H_ sequences and less average support for any given V_L_ compared to V_H_ (Figure 5).

**Figure 5.**
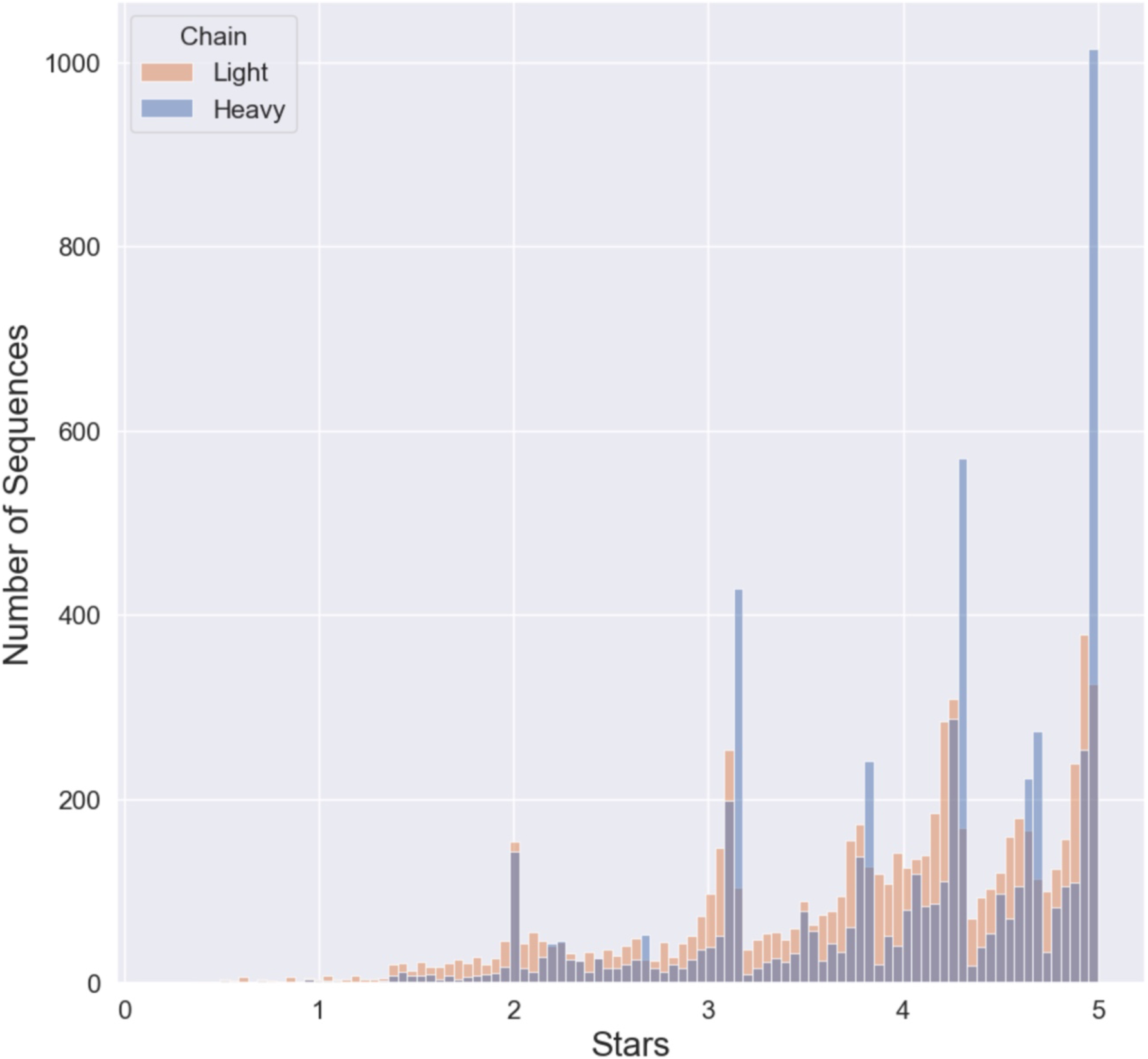
Score distribution of “Stars” awarded to each of the V_L_ and V_H_ sequences reported. Due to the tendency of V_L_ to report more ASVs we see a tendency for a left skewed distribution for V_L_ compared to VH. The sections of high density shown are due to the scoring system which counts the number of matches of biological and technical replicates for a given sequence

### Gibson Assembly-based cloning of recombinant mAbs based on hybridoma sequences

The availability of V_L_ and V_H_ sequences from hybridomas allowed for the conversion of the mAbs they produced into recombinant form. We employed the hybridoma-derived sequences to design V_L_ and V_H_ gene fragments that were used to generate R-mAb mammalian expression plasmids in four fragment Gibson Assembly reactions (Figure 6a).

**Figure 6.**
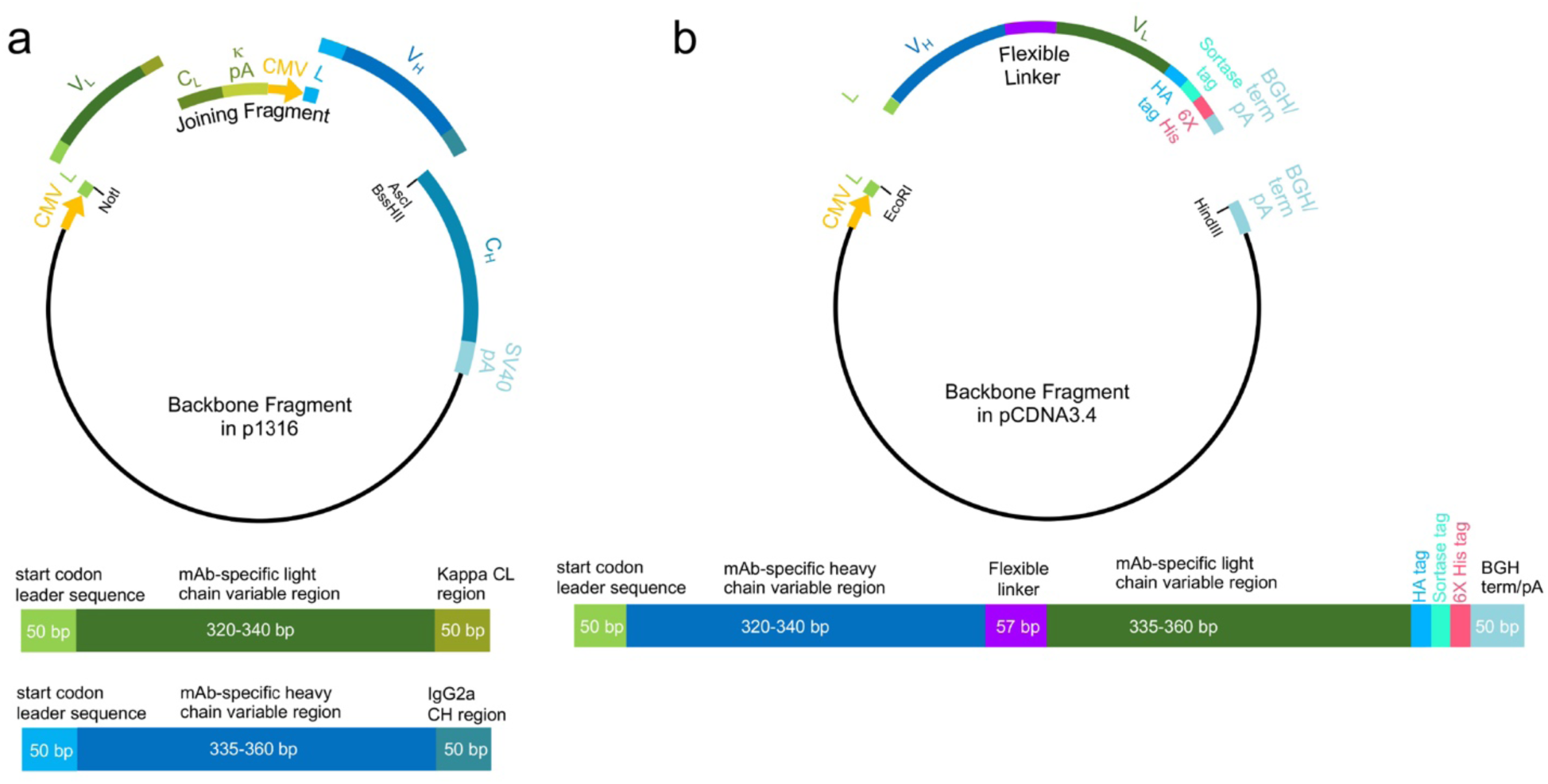
R-mAb and scFv Gibson Assembly cloning strategy. (a) R-mAb cloning strategy showing a schematic of the four-piece Gibson Assembly based construction of the R-mAb expression plasmid and at the bottom a schematic of the V_L_ and V_H_ gene fragments. (b) scFv cloning strategy showing a schematic of the two-piece Gibson Assembly based construction of the scFv expression plasmid and at the bottom a schematic of the V_H_ -linker-V_L_ gene fragment.

Each reaction used two common components, namely the expression vector backbone, and the so-called joining fragment ^9, 28^ (Figure 6a). These two fragments were derived from a modified version ^9^ of the p1316 mouse mAb expression plasmid originally developed by Crosnier, Wright and colleagues ^28^ and based on the pTT3 high level mammalian expression plasmid ^29^. The vector backbone fragment (a 6,702 bp NotI-AscI restriction fragment) contained a CMV promoter followed by the coding sequence of the mouse IgG kappa light chain variable domain leader sequence (accession number A0A0G2JDJ8) prior to the NotI site. It also contained a mouse IgG heavy chain constant region downstream of the AscI site (Figure 6a). We had previously generated a variant of the original p1316 expression plasmid by replacing the mouse IgG1 heavy chain constant region present in p1316 ^28^ with the mouse IgG2a heavy chain constant region (accession number P01863) that we obtained from RT-PCR based cloning from the K28/43 hybridoma cell line ^9^. We initially used this plasmid to express the K28/43 mAb, and then employed this as our primary plasmid for expressing R-mAbs cloned by RT-PCR cloning ^9^. We continued to use this as our primary cloning and expression plasmid as IgG2a mAbs are represented at a much lower proportion in mAb collections than are IgG1 mAbs ^3^, enhancing the value of IgG2a R-mAbs for multiplex labeling ^9^. The second plasmid-derived fragment common to all Gibson Assembly reactions corresponded to the PCR amplified 1,700 bp joining fragment ^28^, comprising the kappa light chain constant region and its associated 3’ untranslated region, a second CMV promoter identical to that in the vector backbone, and a start codon followed by the coding sequence of the mouse IgG variable domain 5’ leader sequence (accession number F6XWB2)(Figure 6a).

Each four fragment Gibson Assembly reaction also included a unique pair of synthesized gene fragments comprising the distinct V_L_ and V_H_ coding regions specific to a particular mAb (Figure 6a). One fragment comprised nucleotide sequences encoding the ANARCI predicted IMGT amino acids 1-127 of the V_L_ region, while the other contained amino acids 1-128 of the V_H_ region, each flanked by 50 bp Gibson Assembly overhang sequences (Figure 6a). We found that a key element for successful four-way Gibson Assembly reactions was the proportion of these four fragments (backbone: 50 ng; joining fragment: 50 ng; V_L_ and V_H_ gene fragments: 0.05 pmol each) that corresponds to a 5:1 molar ratio of each of the other three fragments to the plasmid backbone. The products of the Gibson Assembly reactions were transformed, and selected colonies subjected to colony PCR to amplify the VL-joining fragment-VH region of the plasmid, as we had done previously for screening R-mAb colonies from our PCR-based cloning approach ^9^. We used Sanger sequencing to verify the presence and fidelity of the mAb-specific V_L_ and V_H_ sequences, and sequence-positive plasmids were then used to transfect HEK293-6E cells to produce R-mAb TC supes. These TC supes were subsequently evaluated in immunoassays (IB against brain samples, ICC against transfected cells, and/or IHC on brain sections) in each case performed as a side-by-side comparison with the corresponding conventional mAb TC supernatant (Figure 7).

**Figure 7.**
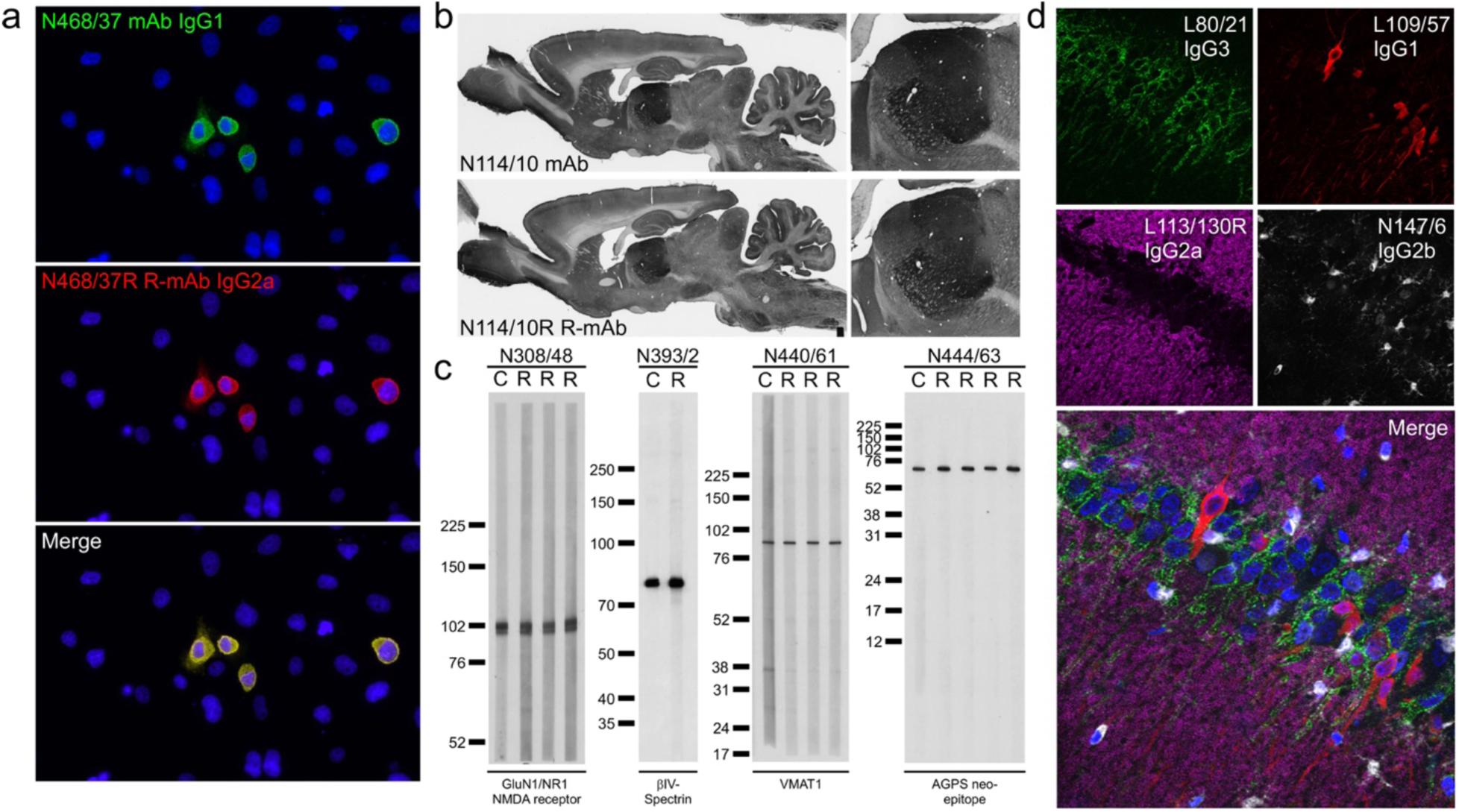
R-mAb evaluation. (a) Immunofluorescence immunocytochemistry on transiently transfected COS-1 cells. Cells were transfected with a plasmid encoding mouse THIK-2 and double immunolabeled with the progenitor N468/37 IgG1 mAb (green) and the N468/37R IgG2a R-mAb (red). Hoechst nuclear labelling is shown in blue. (b) Immunoperoxidase/DAB immunohistochemistry on rat brain sections comparing immunolabeling with the anti-HCN4 hybridoma-generated N114/10 mAb to the N114/10R R-mAb. (c) Strip immunoblots on rat brain samples showing side-by-side comparisons of hybridoma-generated conventional progenitor mAb samples “C” and transfected cell generated R-mAb samples “R”. Numbers to the left of each set denote mobility of molecular weight standards in kD. (d) Immunofluorescence immunohistochemistry on rat brain sections. Multiplex immunolabeling with four mouse mAbs including the subclass-switched R-mAb L113/130R in adult rat hippocampal CA1.

This R-mAb cloning process has been completed employing 410 unique pairs of V_L_ and V_H_ gene fragments in Gibson Assembly reactions. We primarily focused on mAbs whose sequencing yielded only one productive V_L_ and V_H_ domain sequence, and/or which are widely used in the research community based, the number of research publications citing that particular mAb (https://neuromab.ucdavis.edu/publications.cfm). Of these, a very high percentage (381/410 = 93%) yielded at least one plasmid encoding an R-mAb that tested positive in one or more assays in a side-by-side comparison against the progenitor mAb (Figure 7). We also completed this process for eleven mAbs whose hybridomas yielded more than one prominent V_L_ sequence. For ten of these we found one VL-VH combination that yielded a functional mAb, which in six cases was the sequence with the majority of reads, and in four cases the sequence with the minority of reads. One mAb (K60/87) yielded two V_L_ and two V_H_ sequences. In this case all four pairwise combinations were generated of which only one was functional, which corresponded to the V_L_ and V_H_ sequences with the majority of reads. We note that these outcomes are based almost entirely on a single attempt at cloning, expressing, and evaluating each R-mAb using the workflow detailed above. A list of the 392 R-mAbs cloned using this sequence-based approach is provided as Supplementary Table 2. Plasmids for each successfully cloned R-mAb are being deposited at the open access nonprofit plasmid repository Addgene (https://www.addgene.org/), in addition to being registered with and assigned unique RRIDs by The Antibody Registry (https://scicrunch.org/resources).

The availability of the same mAbs in multiple IgG subclasses expands their utility in simultaneous multiplex fluorescent labeling. To enhance the value of our R-mAbs, we generated an IgG2b expression plasmid in which the mouse IgG2a heavy chain constant region of the K89/34 R-mAb was replaced by the corresponding sequence of mouse IgG2b (Uniprot accession number P01867). After functional verification of the expressed subclass-switched K89/34 IgG2b from this plasmid, we used a restriction digest-ligation based cloning approach to transfer the NotI-BssHII fragment of the IgG2a expression plasmids, comprising the VL-joining fragment-VH region of the expression plasmid (Figure 6a), to the NotI-BssHII digested IgG2b expression plasmid backbone. After sequencing and expression, the IgG2b R-mAb TC supes were evaluated in a side-by-side comparison assay against their progenitor mAb or IgG2a R-mAb using subclass-specific secondaries to confirm the subclass switch (Figure 7). To date, we have undertaken the transfer of the V_L_ and V_H_ domains of 85 R-mAbs from the IgG2a plasmid to the IgG2b plasmid with a success rate of 93% (79/85) yielding a functional R-mAb. We have attempted transfer of 53 R-mAbs into a mouse IgG1 expression plasmid with an 81% success rate (43/53). Switching the IgG subclass of R-mAbs allows for multiplex labeling in combinations not possible due to the native IgG subclass of the progenitor mAb (Figure 7). A list of R-mAbs transferred into plasmids encoding alternate heavy chain constant regions is provided as Supplementary Table 3. Plasmids for each of these successfully subclass-switched IgG2b and IgG1 R-mAbs are being deposited at the open access plasmid repository Addgene and registered and assigned unique RRIDs by The Antibody Registry.

### Gibson Assembly-based cloning of scFvs based on hybridoma sequencing

Smaller format/miniaturized recombinant Abs have numerous advantages for immunolabeling including higher resolution imaging and enhanced penetration into samples ^12^. ScFvs that comprise mAb V_L_ and V_H_ domains connected by a flexible linker (Figure 6b) have been in use for over three decades ^10, 11^ as miniaturized derivatives of mAbs. The availability of V_L_ and V_H_ sequences obtained from our high-throughput sequencing allowed us to design gene fragments encoding scFvs. We focused our scFv generation efforts on mAbs that had already been successfully converted to R-mAbs. We used the same VH-linker-VL gene fragment design for each scFv, flanking these sequences with overhangs allowing for Gibson Assembly based cloning into the pCDNA3.1 (or later the pCDNA3.4) mammalian expression plasmid (Figure 6b). In each of the 368 scFv cloning attempts we performed we obtained sequence-positive plasmids. Of these, ≈50% (186/368) yielded at least one plasmid encoding a positive scFv in a transfected cell ICC assay in a side-by-side comparison against the progenitor mAb or R-mAb (Figure 8). We note that these summary numbers are based almost entirely on a single attempt at cloning each scFv using the workflow detailed above. The bulk of these ICC-positive scFvs have been further validated by IHC against brain sections (Figure 8). For fourteen of these attempts for which cloning in the VH-linker-VL design failed to yield a functional scFv, we made a second attempt employing a VL-linker-VH design. Of these, six yielded a functional scFv. A list of the 192 scFvs cloned using this sequence-based approach is provided as Supplementary Table 4. Plasmids for each successfully cloned scFv are being deposited at the open access plasmid repository Addgene and registered with and assigned unique RRIDs by The Antibody Registry.

**Figure 8.**
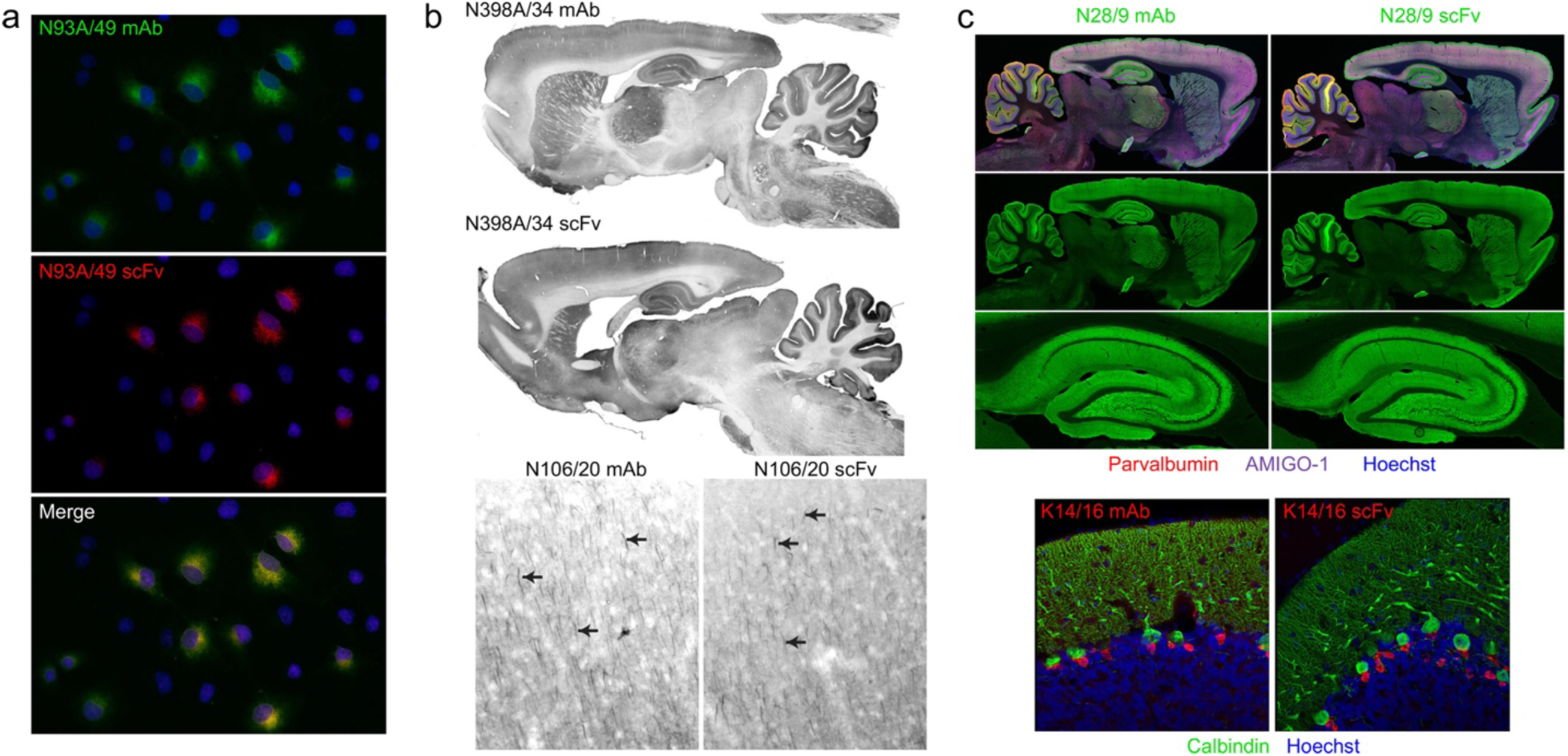
scFv evaluation. (a) Immunofluorescence immunocytochemistry on transiently transfected COS-1 cells. Cells were transfected with a plasmid encoding the mouse GABAB R1 receptor and double immunolabeled with the progenitor N93A/49 mouse mAb (green) and the N93A/49 scFv (red). Hoechst nuclear labelling is shown in blue. (b) Immunoperoxidase/DAB immunohistochemistry on rat brain sagittal sections. The top pair of images show immunolabelling with the anti-GABAA receptor α4 hybridoma-generated N398A/34 mAb (top section) and the N398A/34 scFv (bottom section). The bottom pair of images show immunolabelling in neocortex with the anti-Ankyrin-G hybridoma-generated N106/20 mAb (left section) and the N106/20 scFv (right section). (c) Immunofluorescence immunohistochemistry on rat brain sagittal sections. The top set of images show multiplex immunolabeling of brain (top two rows) and hippocampus (bottom row) with the anti-VGlut1 hybridoma-generated N28/9 mAb (IgG1) (left section) and the N28/9 scFv (right section) in green. The sections were also labelled with the mouse mAbs anti-Parvalbumin L114/3 (IgG2a) in red and the anti-Amigo-1 L86A/37 (IgG2b) in purple. Hoechst nuclear labelling is shown in blue. The bottom pair of images show multiplex immunolabeling of rat cerebellum with the anti-Kv1.2 K^+^ channel hybridoma-generated K14/16 mAb (IgG2b) (left section) and the K14/16 scFv (right section) in green. The sections were also labelled with the mouse anti-Calbindin L109/39 mAb (IgG2b). Hoechst nuclear labelling is shown in blue.

## Discussion

Here we describe a high throughput sequencing-based workflow for determining the sequences of V_L_ and V_H_ domains from cryopreserved hybridomas. This has led to a public sequence database that unequivocally defines at the molecular level a valuable collection of mAbs highly validated for neuroscience research. This workflow includes processing hybridoma samples in 96 well plates to obtain RNA that is used as a template for cDNA synthesis employing a modified 5’-RACE approach. We then use PCR amplification employing non-degenerate semi-nested primers followed by generation of bar-coded sequencing libraries and Illumina sequencing to determine V_L_ and V_H_ domain sequences from hundreds of hybridoma samples in a single sequencing run. A novel bioinformatics platform is then applied to filter and curate the sequences obtained, which are then posted to a public database. The database is accompanied by a variety of useful features for researchers to utilize such as novel quality assessment system, protein annotation, and alignment tools built in across all sequences. Finally, the sequences are used in gene fragment-based cloning of plasmids expressing R-mAbs and scFvs engineered to enhance their utility in multiplex labeling. Together, this represents an effective workflow to preserve in the form of a public database the DNA sequence information defining the mAbs that comprise this large collection of hybridomas, and to use these sequences to develop validated recombinant Ab expression plasmids and make them available through open access non-profit sources.

It is well established that mAbs have tremendous value as renewable research reagents with the potential to endure indefinitely, being produced by immortalized hybridoma cells that can be archived by cryopreservation ^2^. However, it is possible to lose a particular mAb when cryopreserved hybridomas fail to grow when recovered into cell culture, when mAb production is reduced or lost spontaneously, or when mutations arise leading to the production of a mAb with altered properties. In addition, maintenance of cryopreserved hybridoma collections in liquid nitrogen is challenging due to its high cost and labor-intensive nature. Moreover, entire hybridoma collections can be lost upon loss of research funding or the closure of a laboratory. Open access hybridoma banks such as the Developmental Studies Hybridoma Bank at the University of Iowa (https://dshb.biology.uiowa.edu/) represent an important resource in providing longer term preservation of hybridomas above and beyond that afforded by an individual research laboratory. However, they are still subject to the potential loss of hybridoma viability and/or mAb production and fidelity, and the high cost and labor-intensive nature of maintaining cryopreserved hybridomas in liquid nitrogen. Obtaining the sequences of V_L_ and V_H_ domains provides a reliable method for permanently preserving the quintessential identity of a particular mAb and allowing for its functional expression after the hybridoma cells themselves cease to exist. Re-sequencing of extant hybridomas being used for repeated mAb production can also be used as a routine analytical tool to verify that the V_L_ and V_H_ domains remain intact years or decades later after their sequence was originally determined.

We had previously used an RT-PCR based approach employing highly degenerate 5’ PCR primers to generate V_L_ and V_H_ regions amplified from hybridoma-derived cDNA followed by their insertion into plasmids, validation of functional expression and sequencing ^9, 28^. This process is labor intensive and can be prohibitively resource intensive for preserving large hybridoma collections as DNA sequence. Direct sequencing of hybridomas using the approach described here has a higher throughput and requires fewer resources than the cloning based method. Moreover, employing the template switching oligonucleotide based modified 5’-RACE method of cDNA synthesis avoids the problems associated with employing degenerate primers that can cause alterations in the amplified sequences relative to the original sequences present in the source hybridoma due to primer-guided mutations. The workflow we employed can be cost effective when pursued on the scale of 96 well microtiter plates, with the entire process (96 well plate of hybridomas through RNA extraction, cDNA synthesis, PCR amplification and Illumina sequencing) pricing at ≈$20/sample (supply costs plus a generic estimate of $50/hour labor costs). This takes advantage of the high-quality core facilities for RNA purification and DNA sequencing available at UC Davis, although analogous facilities exist at many other academic institutions that could be similarly utilized. Therefore, other users may implement this method to effectively preserve hybridomas in their collection in the form of V_L_ and V_H_ domain DNA sequences.

As a result of our sequencing efforts, we have generated a large public database containing sequences of V_L_ and V_H_ domains of a large collection of highly characterized mAbs. Numerous other public databases of Ab sequences exist, such as the Geneva Antibody Facility ABCD database ^30^, which contains ≈23,000 sequenced Abs against ≈4,000 different targets. These sequences have been mined from publicly available sequences of mAbs of known target specificity. Another extensive collection of Ab sequences is The International Immunogenetics Information System (IMGT) database ^31^. This contains ≈15,000 sequences for rearranged, productive, and presumably functional variable domain mouse IgG heavy and light chains. In this case, sequences are primarily derived from studies of immune repertoire Ab diversity and, for the most part, lack information on the Ab target, although ≈750 sequences in the IMGT database are returned with a search for “hybridoma”. Our public database is distinct from these in that it represents novel sequences of highly characterized mAbs obtained from our in-house sequencing efforts.

The investigation into expression of additional productive (i.e., non-aberrant) V_L_ and/or V_H_ transcripts has revealed a wide range of variation across different monoclonal hybridoma cell lines ^21–27^. This has prompted speculation into the factors contributing to this phenomenon, examples being imperfect allelic exclusion in the splenocyte that gave rise to the hybridoma, or anomalies within the fusion process whereby more than one splenocyte fuses to a single myeloma cell. Compared to a previous analysis of almost 200 rat and mouse hybridoma cell lines ^27^ our sequencing of multiple independent samples from almost 2,000 hybridoma cell lines indicates a lower frequency of multiple productive V_L_ and/or V_H_ transcripts. While we do not know the basis for this, this previous study analyzed both rat and mouse hybridoma cell lines generated in many different laboratories using different approaches and employing different myeloma partners ^27^. Moreover, in this previous study the methods used to obtain V_L_ and V_H_ transcript sequences differed across samples. While a subset was obtained from direct high-throughput sequencing as used here, others came from RT-PCR-based plasmid cloning followed by Sanger sequencing. We used uniform methods to obtain and then process the sequencing data, in the latter case employing a multi-step process that incorporates ASV support, ANARCI prediction, and an in-house scoring system based on reproducibility of obtaining the same sequence from multiple biological and technical replicates. Our dataset also came from a hybridoma collection that, while diverse in the proteins targeted, is otherwise relatively homogenous as it is exclusively mouse hybridomas generated in a single laboratory using the same Sp2/0 myeloma cell line. The one distinction among the hybridomas in our collection was whether they were generated by the original method of PEG/chemical fusion, or the more recent method of electrofusion. However, we found no differences in the incidence of additional among V_L_ and V_H_ transcript sequences among these two sets of hybridomas. We note that the presence of aberrant V_L_ transcripts in Sp2/0-derived hybridoma cells could potentially introduce additional complexity into sample preparation process (e.g., cDNA synthesis and/or RT-PCR amplification steps) and the sequencing data analysis aimed at determining the productive (i.e., non-aberrant) V_L_ transcript repertoire of a given hybridoma cell line. Our curation steps were aimed at addressing these issues to provide a consistent dataset.

Our public database represents a valuable resource for those wishing to use these sequences to recapitulate these mAbs in recombinant form, including those engineered to enhance their utility in multiplex labeling. Cloning of R-mAb expression plasmids employing Gibson Assembly using V_L_ and V_H_ gene fragments designed from hybridoma sequences has advantages over our previous method of RT-PCR based cloning ^9, 28^. First, we avoid the use of degenerate primer sets such as those we ^9^ and many others [e.g., ^24, 28^, etc.] used previously, so that the sequences of the V_L_ and V_H_ domains obtained and used to generate R-mAbs and scFvs are an exact match to the hybridoma. Second, our previous method ^9^ relied on evaluating numerous candidate expression plasmids (sometimes ranging into tens or hundreds) to find those that expressed a functional R-mAb, in part due to the presence of the aberrant V_L_ transcript but also due to mutations that can occur with PCR amplification. Generation of R-mAb expression plasmids based on high quality sequences leads us more directly to a functional R-mAb. We have generated a collection of mouse IgG2a R-mAbs, and in some cases mouse IgG1 and IgG2b versions of the same mAbs, to greatly enhance their utility in combining with other mouse mAbs and/or R-mAbs multiplex fluorescence labeling. We have also generated plasmids encoding functional miniaturized mAbs, in the form of scFvs, that have substantial benefits for immunolabeling due to their small size, such as enhanced sample penetration and increased imaging resolution. Some of these scFvs have already been used to enable a higher resolution correlative light and electron microscopy analysis of cell population within mouse brain ^32^. Moreover, unlike antibodies in their intact IgG format, scFvs can be used as intracellular antibodies or intrabodies ^33^ to effectively report on or manipulate neuronal function ^34^. These publicly posted sequences can be used by researchers to generate a variety of other alternative forms of R-mAbs such as those with heavy chain constant regions from different species, and various fragments, such as Fab, F(ab’)2, Fab2 (monospecific and bi-specific), scFv-Fc, and many others, each of which have distinct properties advantageous to specific applications. The availability of hybridoma-derived sequences also allows for numerous other forms of Ab engineering of recombinant mAbs ^35^ to enhance their binding (affinity and specificity) and biophysical properties (folding and stability), as well as providing insights that could be used in de novo design ^36^. Overall, the workflow described here can be effectively applied to first obtain mAb sequences from hybridomas, and then to use these sequences to generate R-mAbs including those in formats engineered to enhance their utility. In addition, the database and user interface utilized in our study offer a unique and robust platform for standardized data processing. The NeuroMabSeq platform provides enhanced capabilities for data analysis, contributing to a more thorough understanding of the biological processes under investigation.

## Conclusion

We generated an open access publicly available resource in the form of a mAb sequence database and a collection of R-mAbs and scFvs to enable neuroscience research. We first developed a workflow for sequencing and curation of sequences from a large collection of hybridomas. This allowed us to generate a publicly available curated online database of these sequences with numerous attributes to enhance the use of the sequences. We used these sequences to develop a Gibson Assembly-based cloning strategy that led to generation of hundreds of R-mAbs. We also generated additional R-mAb variants with altered IgG subclasses to enhance their utility in multiplex fluorescence labeling. We developed miniaturized versions of these R-mAbs in the form of scFvs to enhance their tissue penetration and imaging resolution due to their small size. All R-mAb and scFv plasmids are publicly available from the open access plasmid resource Addgene.

## Methods

### RNA extraction

RNA was extracted from cryopreserved hybridoma cells in a 96 well plate format. Hybridoma cells maintained in freezing medium (DMEM plus 20% Hyclone Fetal Clone I serum plus 10% DMSO) in liquid nitrogen were rapidly thawed in sets of 12 at 37 °C, and 100 µL aliquots containing ≈0.5-1 x 10^6^ hybridoma cells were immediately transferred to individual wells of a low binding 96 well plate (Costar Catalog# 3896) and stored on ice. Four wells of the plate were kept empty for later inclusion of positive and negative controls.

The plates were centrifuged at 1,600 x g in a swinging bucket microplate rotor (Sorvall Catalog# 75006449H) in a Sorvall Legend T centrifuge (Sorvall Catalog# 75004366) at room temperature. Cell pellets size was estimated by eye, and additional cells were added from the thawed vials on ice if needed, followed by recentrifugation. Supernatant was carefully removed, and pellets were resuspended in 150 µL of PBS. Following centrifugation as above, the PBS wash was removed, and the plate was either stored on ice for use the same day or stored at −80 °C after applying a foil seal (Spectrum Catalog# 634-10749). Hybridoma cryovials containing the remaining cells were also stored at −80 °C as backup for additional RNA extraction at a later date as needed.

Hybridoma cells were lysed by resuspending the cell pellet in 150 µL of RNeasy Lysis Buffer (RLT) (Qiagen Catalog# 1015750) and transferred to the corresponding wells of a 96 well U bottom master block plate (Greiner Catalog# 780215). After sealing with foil (Spectrum Catalog# 634-10749) the plate was vortexed 3 times for 10 s each. The RNA was then purified using a QIAcube HT extraction system (Qiagen Catalog# 9001793) located in the UC Davis School of Veterinary Medicine Real-Time PCR Research and Diagnostics Core Facility (https://pcrlab.vetmed.ucdavis.edu/). The plate of purified RNA (eluted in ≈100 µL molecular biology grade water per well) was foil sealed and stored at −80 °C.

After thawing on ice, each sample was quantified by micro volume spectrometry on a NanoDrop instrument (ThermoFisher Catalog# ND-LITE-PR). After quantification, 10 µL of RNA was transferred to a 96 well hard-shell PCR plate microplate (BioRad Catalog# HSP9601), and the samples in each microplate well normalized to the same concentration by the addition of RNAse-free water. The concentrations ranged from 7-15 ng/µL. Each well within a given plate contained the same RNA concentration to balance the number of sequencing reads for each sample.

### High throughput sequencing of hybridoma V_L_ and V_H_ domains

A schematic of cDNA synthesis and PCR amplification steps is presented as Supplementary Figure 1. All reactions were performed in 96-well and/or 384-well plates and pipetting was carried out with a 96-channel pipette (Gilson Platemaster, Catalog# F110761). Aliquots of RNA from ≈20 of the wells were quantified by fluorometric quantification on a Qubit Flex Fluorometer (ThermoFisher Catalog# Q33327) to confirm the values obtained by Nanodrop. The quality of the RNA in a randomly selected representative number of wells (16 out of 92 wells) was analyzed by microcapillary electrophoresis on a Labchip GX Touch instrument (PerkinElmer Catalog# CLS138162), and plates with RNA samples with a RIN value consistently ≥7 and free of DNA contamination were used for cDNA synthesis. For RNA denaturation, 3 µL of RNA from 4 x 96 well plates was transferred to a 384 well plate with 3 µL of master mix containing 1 µL of a 10 mM stock of dNTPs (NEB Catalog# N0447L; 1.7 mM final concentration) and 0.3 µL of a 10 µM stock of custom IgG chain specific reverse transcription forward primers (0.5 µM final concentration). The oligonucleotide sequences of these constant region chain-specific primers are provided in Supplementary Table 1. RNA was denatured at 72 °C for 3 min and then cooled to 4 °C. After cooling, 4 µL of Reverse Transcription (RT) mix was added to each well. This mix includes 2.5 µL of Template Switching RT Buffer (NEB Catalog# B0466S), 0.5 µL of 25 µM Template Switching Oligo (5 µM final concentration in RT mix; IDT), and 1 µL of Template Switching RT Enzyme Mix (NEB Catalog# M0466L). After mixing, plates were incubated at 42°C for 90 min, and the reaction was stopped by incubating at 85 °C for 5 min. Plates containing reverse transcribed cDNA were stored at 4 °C up to overnight or at −20 °C for longer.

Amplification of V_L_ and V_H_ domain sequences by PCR was performed in a new 384 well plate. cDNA was amplified with a cocktail of four nested constant region chain-specific primers on one end and barcode-indexed primers, targeting the TSO sequence, on the other. The ninety-six unique inline barcode indices identified each well of a source sample plate. The oligonucleotide sequences of the nested constant region chain-specific PCR primers are provided in Supplementary Table 1. The reaction mix contained 2 µL of cDNA, 3 µL of a 2.5 µM stock of bar-coded TSO-specific forward primers (Eurofins; 0.5 µM final concentration), 3 µL of a 2.5 µM stock of IgG chain-specific PCR primers (0.5 µM final concentration), and 8 µL of 2X Phusion polymerase mix (ThermoFisher Catalog# F-565L), for a total volume of 16 µL. After mixing, PCR amplification was performed by 30 cycles of the AMP program comprising thirty cycles of 15 s at 98 °C followed by 30 s at 62 °C and 30 s at 72 °C. Plates were stored at 4 °C up to overnight or at −20 °C for longer. A representative subset of the PCR products (expected size of the amplicons is ≈500-550 bp) was analyzed by microcapillary electrophoresis on a Labchip GX Touch instrument. Barcoded amplicons from each of the four original 96 well plates were pooled separately and cleaned up with 0.6X volume of Kapa Pure Solid Phase Reversible Immobilization (SPRI) beads (Roche Catalog# KK8001), followed by elution in 40 µL of elution buffer. The concentration of DNA in each pool was quantified by Qubit Fluorometric Quantification. Each of the four pools was converted into one Illumina-barcode indexed sequencing library using the ThruPLEX DNA-Seq HV kit (Takara Bio Catalog# R400740), following the manufacturer’s instructions. An aliquot of each library (expected size of the library is ≈650 bp) was analyzed by microcapillary electrophoresis on a Labchip GX Touch instrument. Libraries from up to twelve 96-well plates were sequenced on one MiSeq run (Illumina, San Diego, CA) with paired-end 300 bp sequencing reads (Illumina, San Diego, CA).

### Bioinformatic processing of hybridoma V_L_ and V_H_ domains

The resulting forward and reverse reads were cleaned, joined bioinformatically, and demultiplexed using a custom in house software pipeline. Primer sequence was used to determine heavy or light chain sequence and removed. TSO sequence was identified and removed. Any sequence containing a ‘N’ character was removed from further consideration. Low quality base pairs (<10 q-values) were removed from the 3’ ends, followed by overlapping of paired reads using HTStream (https://github.com/ibest/HTStream, v. 1.2.0-release). Overlapping reads that met a minimum length threshold of 385 bp were then denoised, demultiplexed, and summarized into amplicon sequence variants (ASVs) using the DADA2 algorithm ^14^ and filtered using a custom R script (https://github.com/ucdavis-bioinformatics/NeuroMabSeq/blob/pipeline/02-Results/02-Hybridoma-DADA2-analysis.RMD).

The resulting sequences were then processed through ANARCI using software downloaded from the SAbPred website (https://opig.stats.ox.ac.uk/webapps/sabdab-sabpred/sabpred/) and employing a custom script (https://github.com/ucdavis-bioinformatics/NeuroMabSeq/blob/pipeline/03-annotate-results.py) to confirm classification of the sequence, and to predict amino acid sequence and numbering corresponding to the ImMunoGeneTics (IMGT) convention (https://www.imgt.org/) ^17, 18^. Finally, the resulting sequences were stored in an SQLite database and made available publicly through a website (https://neuromabseq.ucdavis.edu/) built with the Django web framework and hosted on an Amazon Web Services infrastructure for public usage and access. Django is coupled with an object relational mapper so simple structured data can be uploaded, standardized, and further analyzed. Some further analysis supplied by the database infrastructure is direct access to the light chain nucleotide sequence for IMGT amino acids 1-127 as well as the heavy chain nucleotide sequences corresponding to IMGT amino acids 1-128. A breakdown of the LFR1 -LFR4, CDR-L1 - CDR-L3, HFR1 - HFR4, and CDR-H1 - CDR-H3 regions and sequence fragments are also provided.

### Sequence quality assessment and quality control

Sequences with poor annotation via IMGT V_L_ and V_H_ amino acid prediction were removed from the database. This includes sequences with any framework region (FR) or complement determining region (CDR) with zero length. Some basic requirements that need to be met to be deemed a quality annotation include intact FR1 and FR4 regions, a start codon, and the absence of stop codons in the V_L_ and V_H_ coding regions. ANARCI IMGT prediction was also used to further group sequences into bins as some non-VL and non-VH regions were causing unique sequences using DADA2, but the amino acid prediction was the exact same within a given monoclonal hybridoma. This was likely caused by trimming edges based on quality and ‘N’ values. Finally, prior to final scoring, ASV results with < 10% of read support were removed.

Each individual sequence sample was scored to provide users with confidence of sequences in the database. Sequences were assigned a score based on the following heuristic where ASV score can be up to 2 points and match score can be up to 3 points for a total of 5 “stars”:

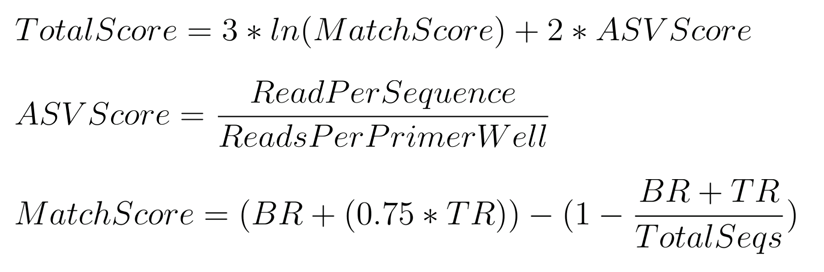

BR refers to biological replicates, while TR refers to technical replicates. In our case, biological replicates are distinct subclones from the same parent, and technical replicates are distinct samples of the same subclone. Both classes of replicates are expected to have an identical V_L_ and V_H_ repertoire. ASV score is the ratio of ASV counts for a given V_L_ or V_H_ sequence over the total ASV count for that sample. For reference, mAb IDs have the following format and scores were assigned in the following manner.

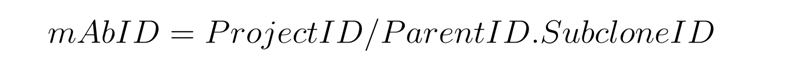

Assigned scores are meant to define the robustness of any particular V_L_ or V_H_ sequence as representing the sequence of the transcripts in that hybridoma cell line. Sequences with a final score of less than 3 were filtered out. Subsequently, the number of sequences returned for a given hybridoma was determined. Sequences were grouped by subclone level name, chain type, plate, sequence, and mAb ID, and the number of occurrences was determined. Data were finally grouped based on the parent hybridoma name, whether it was a V_L_ or V_H_ sequence, and then the assignment of the average number of separate sequences obtained rounded to the nearest integer.

### BLAT feature built into and across the database

BLAST like alignment tool or BLAT ^37^, was incorporated into the database and website to allow for comparison of sequences across parents, subclones, and projects. Inclusion of this tool also allows for insight into sequences and how they might vary across targets, and to perform various forms of quality control. Users can input any nucleotide or amino acid sequence of interest and query the entire database for similar sequences. Search filters can be implemented to refine the search results, and users are immediately linked to the score and accession links for mAbs in the database. Similarly, users can explore the database of nucleotide and amino acid sequences and their quality scores, and within any given entry select the “Blat Sequence” button to cross compare this sequence to all other sequences in the database.

### Generation of R-mAb expression plasmids

Nucleotide sequences corresponding to the ANARCI predicted IMGT amino acids 1-127 of the V_L_ region and amino acids 1-128 of the V_H_ region were used to design gene fragments for use in Gibson Assembly reactions. Gene fragments were designed with 50 bp overhangs corresponding to regions in the P1316 plasmid backbone ^9, 28^ (modified to contain a mouse IgG2a heavy chain) and in the joining fragment region ^9, 28^.

> V_L_ Overhangs
>
> 5’: AGACCCAGGTACTCATGTCCCTGCTGCTCTGCATGTCTGGTGCGGCCGCA
>
> 3’: CGGGCTGATGCTGCACCAACTGTATCCATCTTCCCACCATCCAGTGAGCA
>
> V_H_ Overhangs
>
> 5’: CTGTTCTGCTAGTGGTGCTGCTATTGTTCACGAGTCCAGCCTCAAGCAGT
>
> 3’: GCGCGCCCAACAGCCCCATCGGTCTATCCACTGGCCCCTGTGTGTGGAGA

Commercially generated custom gene fragments (Twist Biosciences) were used in four-piece Gibson Assembly reactions. The expression plasmid backbone was generated by AscI/NotI restriction digestion of the T821a K28/43 IgG2a R-mAb expression plasmid ^9^, available from Addgene (Plasmid# 128618) followed by heat inactivation of the enzymes by incubation at 80°C for 20 min. In some cases, gel isolation of the 6,702 bp fragment was performed. The joining fragment was isolated by PCR amplification of the 1,700 bp region from the beginning of the Kappa light chain constant region to the end of the heavy chain leader sequence, using primers 21 and 26 and using PCR conditions as previously described ^9, 28^.

> Joining fragment PCR primers:
>
> 21: GGGCTGATGCTGCACCAACTGTA
>
> 26: ACTGCTTGAGGCTGGACTCGTGAACAATAGCAGC

After gel purification of the joining fragment, 50 ng (≈ 0.05 pmol) was used in a four-piece Gibson Assembly reaction with 0.05 pmol each of synthetic V_L_ and V_H_ region gene fragments (Twist Biosciences), and 50 ng ≈ 0.01 pmol of the NotI/AscI digested plasmid backbone, corresponding to a 5:1 molar ratio of each of the fragments to the plasmid backbone. Gibson Assembly was performed using NEB HiFi Gibson Assembly Master mix (NEB Catalog# E2621L). Gibson Assembly reaction products were transformed into NEB Turbo high efficiency chemically competent cells (NEB Catalog# C2984I) and plated on LB + Ampicillin plates. Ampicillin-resistant colonies were subjected to colony PCR employing the UpNotI and Rev IgG2a primers that flank the 5’ end of the V_L_ region and the 3’ end of the V_H_ region, respectively.

> UpNotI: 5’-TTTCAGACCCAGGTACTCAT-3’
>
> Rev IgG2a: 5’- ACCCTTGACCAGGCATCCTAGAGT- 3’

Colony PCR products were analyzed on an agarose gel, and colonies yielding an ≈2.4 kB product were isolated from a duplicate patch plate and grown in liquid culture as a source of plasmid DNA (Qiagen QIAprep Spin Miniprep Kit, Catalog# 27106) for sequencing and expression. Sanger sequencing was performed (Quintara Biotech) on an aliquot of the plasmid DNA using sequencing primers allowing for sequencing of the V_L_ and V_H_ regions:

> V_L_: CCTTAGAAGGGAAGATAGGATGG
>
> V_H_: GGATGGTCCACCCAAGAGG

Generation of IgG1 and IgG2b expression plasmids was accomplished by performing NotI/BssHII restriction digests on IgG2a R-mAb expression plasmids generated as described in the preceding section, or in our previous publication ^9^. This digest yields two products, one of ≈2.5 kbp corresponding to the VL-joining fragment-VH region, and the other the 6.7 kbp IgG2a vector backbone. The smaller fragment was gel isolated and used in ligations with gel purified vector backbones prepared from similar restriction digests of analogous IgG1 and IgG2b R-mAb expression plasmids. The K28/43 IgG1 expression plasmid ^9^ derived from the P1316 IgG1 expression plasmid ^28^, was the source of the IgG1 vector backbone. An IgG2b expression plasmid encoding the K89/34 IgG2b R-mAb was generated to order by Genscript (https://www.genscript.com/) by replacing the mouse IgG2a (γ2a) heavy chain constant region of the K89/34 IgG2a R-mAb ^9^ with the corresponding sequence of the mouse IgG2b (γ2b) constant region (Uniprot accession number P01867). Ligations of individual R-mAb VL-joining fragment- V_H_ into one or the other of these vector backbones were transformed into chemically competent XLI-BLUE cells and plated on LB + Ampicillin plates. Ampicillin-resistant colonies were grown in liquid culture as a source of plasmid DNA (Qiagen QIAprep Spin Miniprep Kit, Catalog# 27106) for sequencing and expression. In this case, expression and verification of immunoreactivity was performed first, followed by sequence verification of any subclass-switched R-mAb plasmids whose TC supe exhibited immunoreactivity comparable to the corresponding mAb and/or IgG2a R-mAb by either IB, ICC or IHC.

### scFv cloning

Nucleotide sequences corresponding to the ANARCI predicted IMGT amino acids 1-127 of the V_L_ region and amino acids 1-128 of the V_H_ region were used in combination with a synthetic linker sequence to design an scFv gene fragment for use in Gibson Assembly reactions. The gene fragment encoded a leader-VH-linker-VL scFv followed by HA, sortase and 6xHis tags. The encoded leader amino acid sequence was MGWSCIILFLVATATGVHS, and the encoded linker amino acid sequence was GGGGSGGGGSGGGGSGGGS. Gene fragments were designed with overhangs corresponding to flanking regions in the polylinker cloning region of pCDNA3.4. Gene fragments (Twist Biosciences) were used in two-piece Gibson Assembly reactions. The initial subset of scFvs were cloned into the pcDNA3.1 expression plasmid. The plasmid backbone was generated by EcoRI/HindIII restriction digestion of the N52A/42 scFv originally generated to order by Genscript. The second subset of scFvs were cloned into the pcDNA3.4 expression plasmid. In this case, the plasmid backbone was generated by EcoRI/HindIII restriction digestion of the K89/34 scFv originally generated to order by Genscript. These digestions were followed by heat inactivation of the enzymes by incubation at 80 °C for 20 min. In some cases, gel isolation of the 6,702 bp fragment was performed. The plasmid backbone was then used in a two-piece Gibson Assembly reaction with the synthetic scFv gene fragment (0.05 pMol each) and the digested plasmid backbone (50 ng ≈ 0.01 pmol), corresponding to a 5:1 molar ratio of the scFv gene fragment to the plasmid backbone. Gibson Assembly was performed using NEB HiFi Gibson Assembly Master mix (NEB Catalog# E2621L). The scFv Gibson Assembly reaction products were transformed into NEB Turbo high efficiency chemically competent cells (NEB Catalog# C2984I) and plated on LB + Ampicillin plates. Ampicillin-resistant colonies were selected from transformation plates and liquid bacterial cultures grown as a source of plasmid DNA (Qiagen QIAprep Spin Miniprep Kit, Catalog# 27106) for sequencing and expression. Sanger sequencing was performed (Quintara Biotech) on an aliquot of the plasmid DNA using the CMV-F sequencing primer allowing for sequencing of the scFv coding region:

> CMV-F:5’ - CGCAAATGGGCGGTAGGCGTG - 3’

### R-mAb and scFv expression from transfected mammalian cells

Aliquots of sequence positive R-mAb expression plasmids were used to transfect HEK2936E cells as described, and conditioned media from the transfected cells was harvested 6 days post-transfection ^29^. The conditioned media or R-mAb tissue culture supernatant (R-mAb TC supe) was used for functional testing by immunocytochemistry on transiently transfected cells expressing the target protein, immunoblot against rat brain samples, and/or immunohistochemistry against rat brain sections ^5^.

Aliquots of sequence positive scFv expression plasmids were used to transfect HEK293T cells (ATCC Cat No CRL-3216) using Lipofectamine 2000 (Invitrogen Catalog# 11668019), and conditioned media from the transfected cells was harvested 6 days post-transfection. The conditioned media or scFv TC supe was used for functional testing by immunocytochemistry on transiently transfected cells expressing the target protein and/or immunohistochemistry against rat brain sections ^5^.

### COS cell immunofluorescence immunocytochemistry

R-mAb and scFv TC supes were screened for immunoreactivity in an immunofluorescence assay against transiently transfected COS-1 cells cultured in 96-well plates. COS-1 cells were plated in black, clear bottom 96-well plates (Greiner Catalog# 655090) at a density of 4,700 cells/well. After overnight incubation, each well received 50 ng of plasmid DNA encoding the R-mAb target protein plus Lipofectamine 2000 at a 1:1 ratio as described above. On day 3 post-transfection, cells were washed 3 times with DPBS (138 mM NaCl, 2.67 mM KCl, 1.47 mM KH2PO4, 8.1 mM Na2HPO4, 1 mM CaCl2 and 1 mM MgCl2), pH 7.4 and then fixed using 3.0% formaldehyde (prepared fresh from paraformaldehyde) in DPBS plus 0.1% Triton X-100 on ice for 20 min. Cells were washed 3 times with DPBS/0.1% Triton X-100, and blocked with Blotto/0.1% Triton X-100 for 1 hr. For primary Ab labeling, R-mAb TC supes were used without dilution and hybridoma-generated purified mAb (at 10 µg/mL) or mAb TC supe (at 1:3 dilution) controls were prepared in COS-1 cell culture medium. Primary Abs were incubated at room temperature for 1 hr and cells were washed 3 x 10 min with Blotto/0.1% Triton X-100. Secondary labeling was performed at room temperature for 45 min using subclass-specific, anti-mouse secondary Abs conjugated to Alexa Fluors (ThermoFisher, Catalog#/IgG subclass/Alexa Fluor dye conjugates: (A-21121/IgG1/488 and A-21241/IgG2a/647) and diluted to 1.3 µg/mL in Blotto/0.1% Triton X-100. Hoechst 33342 (ThermoFisher Catalog# H3570) was used at 0.1 µg/mL in the secondary Ab cocktail to stain nuclear DNA. Cells were washed 3 x 10 min with DPBS/0.1% Triton X-100. Imaging was performed using a Zeiss M2 AxioImager microscope. Images were processed using Axiovision (Carl Zeiss Microimaging, RRID:SCR_002677 and Fiji (NIH, RRID:SCR_002285) software.

For higher resolution imaging, COS-1 cells were plated on poly-L-lysine coated #1.5 glass cover slips and cultured overnight followed by transfection with plasmids encoding the target protein. Cells were fixed and immunolabeled as described in the previous section. Images were acquired on a Zeiss AxioImager M2 microscope using a 40x/0.8 NA plan-Apochromat oil-immersion objective and an AxioCam MRm digital camera. Optical sections were acquired using an ApoTome 2 structured illumination system (Carl Zeiss MicroImaging). Imaging and post processing were performed in Axiovision and Photoshop (Adobe Systems; RRID:SCR_014199).

### Immunoblotting against brain samples

R-mAb TC supes were evaluated by immunoblots against rat brain samples. Immunoblots were performed on crude rat brain membrane (RBM) fraction prepared from adult rat brain as previously described ^38^. Following determination of protein concentration by BCA assay (ThermoFisher Catalog# 23227), 3 mg of RBM protein was loaded onto a single well formed by a curtain comb that formed one 12 cm wide well of a large format (18 x 16 cm) SDS-PAGE gel, electrophoresed to size-fractionate the proteins, and transferred onto a nitrocellulose membrane (BioRad Catalog# 1620115). The nitrocellulose membrane was cut into 30 vertical strips (4 mm wide) such that each contained 100 µg of RBM protein. All remaining procedures were performed at RT. Strips were blocked for 1 h in BLOTTO. Primary Ab incubation was performed using undiluted R-mAb TC supes. Comparison strips were incubated in hybridoma generated TC supes (diluted 1:2) or pure mAbs (at 1 µg/mL) diluted in BLOTTO. Following primary Ab incubation, and 3 x 5 min washes in BLOTTO, strips were incubated for 1 h in a volume balanced cocktail of HRP-conjugated affinity-purified goat anti-mouse IgG1, IgG2a and IgG2b secondary Ab (Jackson Immunoresearch Catalog# 115-035-205, 115-035-206 and 115-035-207, respectively) diluted 1:20,000 in BLOTTO. Following 3 x 5 min washes in PBS the chemiluminescent signal was generated by incubation in Western Lightning Plus ECL substrate (ThermoFisher Catalog# 50-904-9325) and subsequently visualized on HyBlot CL film (Denville Scientific Catalog# E3218).

### Immunoperoxidase immunolabeling of free-floating rat brain sections

All experimental procedures were approved by the UC Davis Institutional Animal Care and Use Committee and conform to guidelines established by the National Institutes of Health (NIH). Rats were deeply anesthetized with 60 mg/kg Nembutal sodium solution (pentobarbital, Oak Pharmaceuticals Catalog# 76478-501-20) through intraperitoneal injection, followed by boosts as needed. Once completely anesthetized, rats were transcardially perfused with 25 mL of ice-cold PBS containing 10 U/mL heparin, followed by an ice-cold fixative solution of 4% FA (freshly prepared from PFA, Sigma-Aldrich Catalog# 158127) in 0.1 M sodium phosphate buffer (PB), pH 7.4, using a volume of 0.5 mL fixative solution per gram of rat weight. Following perfusions, brains were removed from the skull and cryoprotected in 10% sucrose in 0.1 M PB for 30 minutes at RT, then transferred to a solution of 30% sucrose in 0.1 M PB for 24-48 h, until they equilibrated. Following cryoprotection, all brains were frozen, and sagittal sections were cut at 30 µm thickness on a freezing stage sliding microtome (Richard Allen Scientific). Sections were collected in 0.1 M PB containing 10 mM sodium azide and processed for free floating immunohistochemistry. Immunoperoxidase immunolabeling of brain sections with R-mAb or scFv TC supes and corresponding hybridoma-generated mAb TC supes or pure mAbs was performed essentially as previously described ^39^. All incubations and washes were at RT with slow agitation, unless stated otherwise. Briefly, free-floating sections were washed 3 x 5 min in 0.1 M PB. Sections were incubated in blocking buffer (10% goat serum in 0.1 M PB, 0.3% Triton X-100, and 10 mM sodium azide) for 1 h. Immediately after blocking, sections were placed into undiluted R-mAb or scFv TC supes supplemented with goat serum (final 5% v/v) and Triton X-100 (final 0.3%) or with mAb TC supes or pure mAb preparations diluted in blocking buffer, followed by overnight incubation at 4 °C with slow agitation. Sections were then washed 3 x 10 min in 0.1 M PB. For R-mAb and mAb primary antibodies, the sections were incubated for 1 h with a volume-balanced cocktail of biotinylated goat anti-mouse IgG subclass-specific secondary antibodies (Jackson Immunochemicals Catalog# 115-065-205 anti-IgG1, Catalog# 115-065-206 anti-IgG2a, and Catalog# 115-065-207 anti-IgG2b) diluted 1:2000 in blocking buffer. For scFvs, sections were incubated for 1 h in anti-HA 2-2.2.14 IgG1 mouse mAb (ThermoFisher Catalog# 26183; RRID:AB_10978021) diluted 1:2000 and following 3 x 10 min washes in 0.1 M PB, sections were incubated for 1 h in Biotin SP−conjugated goat anti-mouse IgG1 secondary Ab (Jackson Immunochemicals Catalog# 115-065-205). For all samples, after 3 x 10 min washes in 0.1 M PB, sections were incubated for 1 h in ABC solution (Peroxidase Standard Vectastain Elite ABC-HRP Kit, Vector Laboratories, Catalog# PK-6100). After 2 x 5min washes in 50 mM Tris-HCl, pH 7.5, sections were incubated in a substrate solution containing 0.4 mg/mL 3,3-diaminobenzidine (Millipore/Sigma Catalog# D5905), 3 mg/mL nickel ammonium sulfate hexahydride (ThermoFisher Catalog# N48) plus 0.003% hydrogen peroxide (Millipore/Sigma Catalog# HX0635) until sufficient reaction product was obtained. Sections were transferred to 50 mM Tris-HCl, pH 7.5 to stop development, rinsed two times with 50 mM Tris-HCl, pH 7.5, and mounted on Superfrost Plus (Fisher Scientific Catalog# 12-550-15) microscope slides. Mounted sections were dehydrated successively in 70%, 95%, and 100% (twice) ethanol, for 5 min at RT each. Sections were cleared in Citrus Clearing Solvent (Richard-Allan Scientific/ThermoFisher Catalog# 8301) two times for 10 min each, and cover slipped with DEPEX mountant (Electron Microscopy Sciences Catalog# 13515). The slides were analyzed and imaged after curing overnight in a fume hood.

### Multiplex immunofluorescence labeling of brain sections

Multiplex immunofluorescence labeling of rat brain sections was performed essentially as described previously ^3, 40^. Rats were perfused, brains removed, cryoprotected, frozen and sectioned as above. Sections were collected in 0.1 M PB and processed immediately for immunohistochemistry. Free-floating brain sections were blocked with 10% goat serum in 0.1 M PB containing 0.3% Triton X-100 (vehicle) for 1 h at RT and then incubated overnight at 4°C in vehicle containing different combinations of primary antibodies. HA-tagged scFvs were detected using mouse mAb 2-2.2.14 (ThermoFisher Catalog# P126183, RRID:AB_10978021). The following day sections were washed 4 x 5 min with vehicle, and then incubated for 1 h at RT in mouse IgG subclass-specific secondary Abs as described previously ^3, 41^. Sections were then washed 2 x 5 min with 0.1 M PB, 2 x 5 min with 0.05 M PB and mounted on Superfrost Plus (ThermoFisher Catalog# 12-550-15) microscope slides, air dried and cover slipped with ProLong Gold Antifade Mountant (ThermoFisher Catalog# P36930). Images were obtained on a ZeissObserver.Z1 microscope with Apotome 2.0. Imaging and post-imaging processing was performed in Zen Blue (Zeiss, RRID:SCR_013672) and Adobe Photoshop software, taking care to maintain any linear differences in signal intensities present in the original samples.

### Software Availability

The analysis pipeline and website are publicly available at

https://github.com/ucdavis-bioinformatics/NeuroMabSeq/tree/website_2.0

https://github.com/ucdavis-bioinformatics/NeuroMabSeq/tree/pipeline

## Acknowledgments

We thank Jessica Bernal, Camelia Dumitras, Henrique Noro Frizzo, Quynh Giang Le, Natasha Mariano, Sara McGrath, Phillip Singh, Ebun Smith and Deborah van der List for their expert technical contributions, and Dr. JoAnne Engebrecht for helpful advice on the project.

## Author contributions

KGM: Conceptualization, methodology, formal analysis, investigation, writing—original draft preparation, writing—review and editing, visualization

BG: Conceptualization, methodology, investigation, writing—review and editing

SH: Conceptualization, methodology, investigation, formal analysis, writing—review and editing DB-W: Conceptualization, methodology, investigation, writing—review and editing

CG-O: Conceptualization, methodology, investigation, writing—review and editing

KMT: Conceptualization, methodology, investigation, writing—review and editing

MEG: Conceptualization, methodology, investigation, writing—review and editing

MB: Conceptualization, methodology, investigation, writing—review and editing

LM: Conceptualization, methodology, investigation, writing—review and editing

MLS: Conceptualization, supervision

LF: Conceptualization, writing—original draft preparation, writing—review and editing, supervision

JST: Conceptualization, writing—original draft preparation, writing—review and editing, visualization, supervision, project administration, funding acquisition

All authors have read and agreed to the published version of the manuscript.

