## Supplementary Figure 1 for "NeuroMabSeq: high volume acquisition, processing, and curation of hybridoma sequences and their use in generating recombinant monoclonal antibodies and scFvs for neuroscience research"

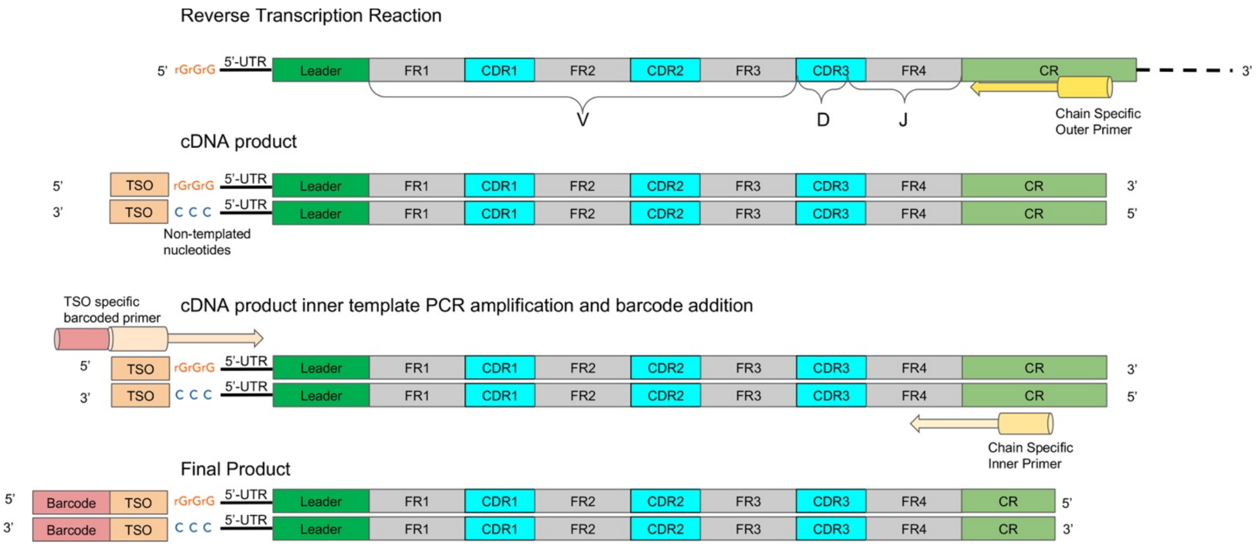


**Supplementary Figure 1**. cDNA synthesis and PCR amplification overview. Schematic shows location of nested chain-specific primers used for cDNA synthesis (outer) and PCR amplification (inner). Also shown is the location of the template switching oligonucleotide. Schematic depicts the synthesis and amplification of VH domain cDNA, analogous steps are used for VL domains. FR: framework regions; CDR: complementarity determining regions; CR: constant region.
