## Supplementary Table 1 for "NeuroMabSeq: high volume acquisition, processing, and curation of hybridoma sequences and their use in generating recombinant monoclonal antibodies and scFvs for neuroscience research"

**Supplementary Table 1. List of primers and barcodes.**

| Oligonucleotide | Vendor | Cat # | Sequence |
|---|---|---|---|
| RT Chain Specific Outer Primer mix | | | |
| RT-mIGK | IDT | 271038654 | TTGTCGTTCACTGCCATCAAT*C |
| RT-mIGHG | IDT | 271038655 | AGCTGGGAAGGTGTGCACA*C |
| RT-mIGL | IDT | 271038656 | GGGGTACCATCTACCTTCCA*G |
| RT-mIGHG3-NEW | IDT | 271038657 | GTACAGATGAGACTGTGCGCACA*C |
| TSO-short | IDT | 408473058 | /5Biosg/AAGCAGTGGTATCAACGCAGAGTACA TrGrGrG |
| PCR Chain Specific Reverse Inner Primer Mix | | | |
| 10for-mIgh3 | IDT | 270929982 | GGATAGACAGATGGGGCTGTTGTTGTAG*C |
| PCR-R-mIGL | IDT | 270929983 | ATCGTACACACCAGTGTGG*C |
| 6-REV-LC | IDT | 270929984 | GGATACAGTTGGTGCAGCAT*C |
| 10-REV-HC1 | IDT | 270929985 | ATAGACAGATGGGGGTGTCGTTTTGG*C |

| F Primer, ISPCR (Barcoded) PCR-SMART indices | Sequence |
|---|---|
| 01-SMARTindex | CAGCGTCAGTGGTATCAACGCAGAGTACA |
| 02-SMARTindex | GATCACCAGTGGTATCAACGCAGAGTACA |
| 03-SMARTindex | ACCAGTCAGTGGTATCAACGCAGAGTACA |
| 04-SMARTindex | TGCACGCAGTGGTATCAACGCAGAGTACA |
| 05-SMARTindex | ACATTACAGTGGTATCAACGCAGAGTACA |
| 06-SMARTindex | GTGTAGCAGTGGTATCAACGCAGAGTACA |
| 07-SMARTindex | CTAGTCCAGTGGTATCAACGCAGAGTACA |
| 08-SMARTindex | TGTGCACAGTGGTATCAACGCAGAGTACA |
| 09-SMARTindex | TCAGGAACAGTGGTATCAACGCAGAGTACA |
| 10-SMARTindex | CGGTTAACAGTGGTATCAACGCAGAGTACA |
| 11-SMARTindex | TTAACTACAGTGGTATCAACGCAGAGTACA |
| 12-SMARTindex | ATGAACACAGTGGTATCAACGCAGAGTACA |
| 13-SMARTindex | CCTAAGACAGTGGTATCAACGCAGAGTACA |
| 14-SMARTindex | AATCCGACAGTGGTATCAACGCAGAGTACA |
| 15-SMARTindex | GGCTGCACAGTGGTATCAACGCAGAGTACA |
| 16-SMARTindex | TACCTTACAGTGGTATCAACGCAGAGTACA |
| 17-SMARTindex | TCTTAATCAGTGGTATCAACGCAGAGTACA |
| 18-SMARTindex | GTCAGGTCAGTGGTATCAACGCAGAGTACA |
| 19-SMARTindex | ATACTGTCAGTGGTATCAACGCAGAGTACA |
| 20-SMARTindex | TATGTCTCAGTGGTATCAACGCAGAGTACA |
| 21-SMARTindex | GAGTCCTCAGTGGTATCAACGCAGAGTACA |
| 22-SMARTindex | GGAGGTTCAGTGGTATCAACGCAGAGTACA |
| 23-SMARTindex | CACACTTCAGTGGTATCAACGCAGAGTACA |
| 24-SMARTindex | CCGCAATCAGTGGTATCAACGCAGAGTACA |
| 25-SMARTindex | TTTATGCAGTGGTATCAACGCAGAGTACA |
| 26-SMARTindex | AACGCCCAGTGGTATCAACGCAGAGTACA |
| 27-SMARTindex | CAAGCACAGTGGTATCAACGCAGAGTACA |
| 28-SMARTindex | GCTCGACAGTGGTATCAACGCAGAGTACA |
| 29-SMARTindex | GCGAATCAGTGGTATCAACGCAGAGTACA |
| 30-SMARTindex | TGGATTCAGTGGTATCAACGCAGAGTACA |
| 31-SMARTindex | ACCTACCAGTGGTATCAACGCAGAGTACA |
| 32-SMARTindex | CGAAGGCAGTGGTATCAACGCAGAGTACA |
| 33-SMARTindex | AGATAGAACAGTGGTATCAACGCAGAGTACA |
| 34-SMARTindex | TTGGTAAACAGTGGTATCAACGCAGAGTACA |

| | |
|---|---|
| 35-SMARTindex | GTTACCAACAGTGGTATCAACGCAGAGTACA |
| 36-SMARTindex | CGCAACAACAGTGGTATCAACGCAGAGTACA |
| 37-SMARTindex | TGGCGAAACAGTGGTATCAACGCAGAGTACA |
| 38-SMARTindex | ACCGTGAACAGTGGTATCAACGCAGAGTACA |
| 39-SMARTindex | CAACAGAACAGTGGTATCAACGCAGAGTACA |
| 40-SMARTindex | GATTGTAACAGTGGTATCAACGCAGAGTACA |
| 41-SMARTindex | CTCTCGATCAGTGGTATCAACGCAGAGTACA |
| 42-SMARTindex | TGACACATCAGTGGTATCAACGCAGAGTACA |
| 43-SMARTindex | AAGACAATCAGTGGTATCAACGCAGAGTACA |
| 44-SMARTindex | ACAGATATCAGTGGTATCAACGCAGAGTACA |
| 45-SMARTindex | TAGGCTATCAGTGGTATCAACGCAGAGTACA |
| 46-SMARTindex | CTCCATATCAGTGGTATCAACGCAGAGTACA |
| 47-SMARTindex | GCATGGATCAGTGGTATCAACGCAGAGTACA |
| 48-SMARTindex | AATAGCATCAGTGGTATCAACGCAGAGTACA |
| 49-SMARTindex | GTGCCATACAGTGGTATCAACGCAGAGTACA |
| 50-SMARTindex | TCGAGGTACAGTGGTATCAACGCAGAGTACA |
| 51-SMARTindex | CACTAATACAGTGGTATCAACGCAGAGTACA |
| 52-SMARTindex | GGTATATACAGTGGTATCAACGCAGAGTACA |
| 53-SMARTindex | CGCCTGTACAGTGGTATCAACGCAGAGTACA |
| 54-SMARTindex | AATGAATACAGTGGTATCAACGCAGAGTACA |
| 55-SMARTindex | ACAACGTACAGTGGTATCAACGCAGAGTACA |
| 56-SMARTindex | ATATCCTACAGTGGTATCAACGCAGAGTACA |
| 57-SMARTindex | AGTACTCAGTGGTATCAACGCAGAGTACA |
| 58-SMARTindex | ATAAGACAGTGGTATCAACGCAGAGTACA |
| 59-SMARTindex | GGTGAGCAGTGGTATCAACGCAGAGTACA |
| 60-SMARTindex | TTCCGCCAGTGGTATCAACGCAGAGTACA |
| 61-SMARTindex | GAAGTGCAGTGGTATCAACGCAGAGTACA |
| 62-SMARTindex | CAATGCCAGTGGTATCAACGCAGAGTACA |
| 63-SMARTindex | ACGTCTCAGTGGTATCAACGCAGAGTACA |
| 64-SMARTindex | CAGGACCAGTGGTATCAACGCAGAGTACA |
| 65-SMARTindex | AAGCTCCAGTGGTATCAACGCAGAGTACA |
| 66-SMARTindex | GACGATCAGTGGTATCAACGCAGAGTACA |
| 67-SMARTindex | TCGTTCCAGTGGTATCAACGCAGAGTACA |
| 68-SMARTindex | CCAATTCAGTGGTATCAACGCAGAGTACA |
| 69-SMARTindex | AGTTGACAGTGGTATCAACGCAGAGTACA |
| 70-SMARTindex | AACCGACAGTGGTATCAACGCAGAGTACA |
| 71-SMARTindex | CAGATGCAGTGGTATCAACGCAGAGTACA |
| 72-SMARTindex | GTAGAACAGTGGTATCAACGCAGAGTACA |
| 73-SMARTindex | GACATCACAGTGGTATCAACGCAGAGTACA |
| 74-SMARTindex | CGATCTACAGTGGTATCAACGCAGAGTACA |
| 75-SMARTindex | CGTCGCACAGTGGTATCAACGCAGAGTACA |
| 76-SMARTindex | ATGGCGACAGTGGTATCAACGCAGAGTACA |
| 77-SMARTindex | ATTGGTACAGTGGTATCAACGCAGAGTACA |
| 78-SMARTindex | GCCACAACAGTGGTATCAACGCAGAGTACA |
| 79-SMARTindex | CATCTAACAGTGGTATCAACGCAGAGTACA |
| 80-SMARTindex | AACAAGACAGTGGTATCAACGCAGAGTACA |
| 81-SMARTindex | GCAGCCTCAGTGGTATCAACGCAGAGTACA |
| 82-SMARTindex | ACTCTTTCAGTGGTATCAACGCAGAGTACA |
| 83-SMARTindex | TGCTATTCAGTGGTATCAACGCAGAGTACA |
| 84-SMARTindex | AAGTGGTCAGTGGTATCAACGCAGAGTACA |
| 85-SMARTindex | CTCATATCAGTGGTATCAACGCAGAGTACA |
| 86-SMARTindex | CCGACCTCAGTGGTATCAACGCAGAGTACA |
| 87-SMARTindex | GGCCAATCAGTGGTATCAACGCAGAGTACA |

| | |
|---|---|
| 88-SMARTindex | AGACCATCAGTGGTATCAACGCAGAGTACA |
| 89-SMARTindex | CGCGGACAGTGGTATCAACGCAGAGTACA |
| 90-SMARTindex | CCTGCTCAGTGGTATCAACGCAGAGTACA |
| 91-SMARTindex | GCGCTGCAGTGGTATCAACGCAGAGTACA |
| 92-SMARTindex | GAACCTCAGTGGTATCAACGCAGAGTACA |
| 93-SMARTindex | TTCGAGCAGTGGTATCAACGCAGAGTACA |
| 94-SMARTindex | AGAATCCAGTGGTATCAACGCAGAGTACA |
| 95-SMARTindex | AGGCATCAGTGGTATCAACGCAGAGTACA |
| 96-SMARTindex | ACACGCCAGTGGTATCAACGCAGAGTACA |

**Supplementary Table 1.** Top table. Sequences of oligonucleotides used for priming cDNA synthesis and for amplification of $V_L$ and $V_H$ domains. Bottom table. Sequences of oligonucleotides used to barcode samples.
