## Supplementary Table 2 for "NeuroMabSeq: high volume acquisition, processing, and curation of hybridoma sequences and their use in generating recombinant monoclonal antibodies and scFvs for neuroscience research"

### Supplementary Table 2. List of R-mAbs successfully cloned via sequence-based cloning.

| R-mAb | Target | R-mAb | Target | R-mAb | Target | R-mAb | Target |
|---|---|---|---|---|---|---|---|
| 1D8 | LRRK2/Dardarin-pSer910 | L114/3 | Parvalbumin | N133/35 | RGS14 | N358/68 | 5-formylcytidine |
| 1F1 | TrpC1 | L114/38 | Parvalbumin | N134/12 | Nav1.8 Na+ channel | N360A/24 | ATF4 |
| 62-3G1 | GABA(A)R, Beta2/3 | L114/81 | Parvalbumin | N138/6 | LRRK2/Dardarin, N-terminus | N363/71 | Kir6.2 K+ channel |
| 8G10 | LRRK2/Dardarin, N-terminus | L115/13 | NPY Neuropeptide Y | N140A/12 | SALM5/LRFN5 | N364/10 | Bral1 |
| A12/18 | Pan-Neurofascin (extracellular) | L117/1 | IRSp53/BAIAP2 | N141/21 | HCN3 | N364/42 | Bral1 |
| D4/11 | Kv2.1 K+ channel | L117/52 | IRSp53/BAIAP2 | N141/28 | HCN3 | N366/60 | Kir6.1 K+ channel |
| D4/154 | Kv2.1 K+ channel | L118/14 | VGAT | N142/28 | GRK6A/B | N367/51 | Shank1 and Shank3 |
| D4/40 | Kv2.1 K+ channel | L118/80 | VGAT | N143/36 | CDY1/2/L1/L2 | N367/62 | Shank3 |
| K14/39 | Kv1.2 K+ channel | L118/135 | VGAT | N143/38 | CDY1/2 | N372A/1 | Kv2.2 K+ channel |
| K19/46 | Kv1.6 K+ channel | L119/106 | NPY2R Neuropeptide Y receptor type 2 | N144/14 | 6xHis | N372B/1 | Kv2.2 K+ channel |
| K28/25 | PSD-95 + PSD-93 | L119/134* | NPY2R Neuropeptide Y receptor type 2 | N144/17 | Gamma-protocadherin-A3 | N372B/60 | Kv2.2 K+ channel |
| K28/27 | PSD-95 | L120/12 | Collybistin | N144/32 | Pan-Gamma-protocadherin-A | N372C/51 | Kv2.2 K+ channel |
| K28/38 | PSD-95 | L120/30 | Collybistin | N145/20 | Pan-GRK | N374/48 | TASK1 K+ channel |
| K28/42 | PSD-95 + PSD-93 | L121/25 | nNOS/NOS1 | N148/30 | Gamma-protocadherin-B2 | N375/67 | Kv3.3 K+ channel |
| K28/58 | PSD-95 | L121/42 | nNOS/NOS1 | N152B/23 | VDAC1 | N377/20 | MAP3K12 |
| K28/74 | PSD-95 | L121/133 | nNOS/NOS1 | N154/32 | JMJD2A | N377/83 | MAP3K12 |
| K28/86 | Pan-MAGUK | L121/136 | nNOS/NOS1 | N155/9 | Uncx | N37A/10 | Kv7.1/KCNQ1 K+ channel |
| K28/94 | PSD-95 | L122/6 | Calretinin | N158/28 | Histone H3.3 | N380/87 | CLN3 |
| K40/17 | Pan-Kvbeta K+ channel | L122/68 | Calretinin | N159/5 | Pan-Gamma-protocadherin-Constant | N382/14 | MFF |
| K47/9 | Kvbeta1.2/KCNAB1 K+ channel | L122/144 | Calretinin | N160/21 | SynDIG3/Tmem91 | N382/69 | MFF (exon 1) |
| K47/42 | Kvbeta1.2/KCNAB1 K+ channel | L123/46 | GABA(A)R, Theta | N161/20 | NKCC1 | N384/63 | Miro2 |
| K55/7* | KChIP1 K+ channel | L124/59 | Bassoon | N165/38 | LAR/PTPRF | N385/21 | Beta1-spectrin |
| K55/29 | KChIP1 K+ channel | L125/121 | Synapsin-3 | N165/43 | Pan-PTPR | N388A/27 | Ankyrin-R/G |
| K55/82 | Pan-KChIP K+ channel | L125/129 | Pan-Synapsin | N166A/26 | OCRL/INPP5b | N389/9 | TRAK1 |
| K56A/50 | CASK | L125/76 | SAPAP3 | N170A/26 | Neurexin-1-Beta | N390/43 | TRAK2 |
| K56A/57 | CASK | L126/93 | SAPAP3 | N173B/13 | Tafazzin | N391/68 | WNK1 |
| K57/27 | Kv4.2 K+ channel (external) | L127/8 | GAD67 | N180/41 | EAAC1 | N393/2 | Beta4-spectrin |
| K57/41 | Kv4.2 K+ channel (external) | L127/12 | GAD65/67 | N183/15 | QKI-7 | N393/76 | Beta4-spectrin |
| K60/73 | KChIP2b K+ channel | L130/1 | Tiam1 | N185/7 | NALCN | N395/68 | Navbeta2 Na+ channel |
| K60/87** | KChIP2b K+ channel | L131/17 | SPHKAP | N186/29 | Dopamine D2 receptor | N396/29 | Navbeta3 Na+ channel |
| K65/35 | CASPR/Neurexin IV | L131/20 | SPHKAP | N191/7 | Frataxin | N397/19 | Nav1.5 Na+ channel |
| K65A/2 | CASPR/Neurexin IV | L131/27 | SPHKAP | N192/12 | Gs protein, alpha subunit | N398A/34 | GABA(A)R, Alpha4 |
| K66/27 | KChIP3 K+ channel | L132/18 | Cav1.2 Ca2+ channel pS1928 | N194/11 | QKI-7b | N399/19 | GABA(A)R, Alpha2 |
| K66/59 | KChIP3 K+ channel | L132/37 | Cav1.2 Ca2+ channel pS1928 | N202/7 | Fig4/Sac3 | N400/24 | CLN6 |
| K67/11 | CASPR2 | N1/12 | KCC2 | N206A/8 | GFAP | N402/13 | Thyroid hormone receptor beta1 |

| | | | | | | | |
|---|---|---|---|---|---|---|---|
| K67/25 | CASPR2 | N4/15 | PINK1 | N207/27 | LRP4 (extracellular) | N402/46 | Thyroid hormone receptor beta1 |
| K69/3* | Nav1.2 Na+ channel | N4/49 | PINK1 | N209C/35 | LRRTM2 | N403/63 | Pan-Thyroid hormone receptor |
| K69/33 | Nav1.2 Na+ channel | N6/38 | VAChT | N210/5 | PhyH/PAHX | N405/74 | Navbeta1 Na+ channel |
| K73/20 | Contactin/F3 | N10/7 | Cavbeta4 Ca2+ channel | N212/7 | TRIP8b (constant) | N406/47 | Clcn5 |
| K74/53 | Nav1.1 Na+ channel | N15/4 | TrpV3 | N212A/34 | TRIP8b (exon 1b) | N408/79 | Npas4 |
| K75/18 | Kv4.3 K+ channel | N15/39 | TrpV3 | N219/5 | RBM17/SPF45 | N410/17 | Kv3.2 K+ channel |
| K75/30 | Kv4.3 K+ channel | N18/28 | PSD-93/Chapsyn-110 | N221/12 | TrpV1 | N411/51 | Arx |
| K78/29 | SK2 K+ channel | N18/30 | PSD-93/Chapsyn-110 | N221/17 | TrpV1 | N414/25 | NCKX4 |
| K87A/10 | Nav1.6 Na+ channel | N19/2 | SAP102 | N225A/10 | FGF13/FHF2, B isoform | N416/57 | GluN3A/NR3A glutamate receptor |
| K89/41 | Kv2.1 K+ channel | N22/21 | Shank1 | N227/21 | MECP2 | N420/24 | EAAC1 |
| K96/7 | Pancortin | N23B/6 | Shank2 | N228A/16 | Lhx6.1 | N421A/85 | ANO5/TMEM16E |
| L6/48 | Slo1/BKAlpha maxi-K+ channel | N25/35 | Kir2.3 K+ channel | N229A/32 | GABA(A)R, Alpha6 | N422/18 | GluN2C/NR2C glutamate receptor |
| L6/60 | Slo1/BKAlpha maxi-K+ channel | N26A/23 | Kv7.2/KCNQ2 K+ channel | N231B/34 | LRRK2/Dardarin, N3 (non-mouse-reactive) | N423/75 | KChIP4 K+ channel |
| L21/32 | GluA2/GluR2 glutamate receptor | N38/8 | Cav1.3 Ca2+ channel | N232/9 | PEX7 | N424/45 | Glycine receptor Alpha3 |
| L23/27 | Kv1.3 K+ channel | N39B/8 | GIT1 | N233/8 | PEX6 | N424/48* | Glycine receptor Alpha3L |
| L24/1 | IP3 receptor, type 1 | N43/6 | Kv7.4/KCNQ4 K+ channel | N238/29 | SAPAP1 | N425/45 | VAChT |
| L24/11 | IP3 receptor, type 1 | N46/30 | ADAM22 (cytoplasmic) | N238/30 | SAPAP1 | N428/12 | WRN |
| L24/18 | IP3 receptor, type 1 | N49A/21 | NGL-1/LRRC4C | N238/31 | SAPAP1/2 | N429/19 | ANO6/TMEM16F |
| L24/19 | IP3 receptor, type 1 | N50/36 | NGL-2/LRRC4 | N241A/34 | LRRK2/Dardarin, C-terminus | N431/64 | GABA(A)R, Pi |
| L24/21 | IP3 receptor, type 1 | N52B/27 | SALM2/LRFN1 | N241A/72 | LRRK2/Dardarin, C-terminus | N432/21 | VSP |
| L28/36 | Kv4.2 K+ channel | N53/32 | BKbeta2 K+ channel | N245/1 | TARPGamma2/Stargazin | N432/63 | VSP |
| L45/30 | SynCAM1 | N55/10 | Cav3.2 Ca2+ channel | N245/36 | TARPGamma2/4/8 | N440/21 | VMAT1 |
| L48A/9 | Cav1.3 Ca2+ channel | N56/9 | S-tag | N245/44 | Thioredoxin | N440/61 | VMAT1 |
| L48A/29 | Cav1.3 Ca2+ channel | N56/21 | FGF14/FHF4 | N250/21 | DNAH7 | N441/35 | ADAM11 |
| L48A/31 | Cav1.3 Ca2+ channel | N57/2 | ADAM22 (extracellular) | N251/14 | DNAH1 | N442/28 | Alg13 |
| L57/23 | Cav1.2/1.3 Ca2+ channel | N59/20 | GluN2B/NR2B glutamate receptor | N253/32 | Notch1 | N444/63 | AGPS neo-epitope |
| L57/46 | Cav1.2/1.3 Ca2+ channel | N64A/36 | TrpC7 | N255/38 | Nav1.4 Na+ channel | N445/27 | TRAAK K+ channel |
| L57/47 | Cav1.2/1.3 Ca2+ channel | N67/15 | TrpC5 | N263/31 | Cav1.2 Ca2+ channel | N446/80 | CELF4/BRUNOL4 |
| L58A/6 | Kv2.1 K+ channel | N68/6 | Nav1.7 Na+ channel | N270/47 | Synaptotagmin-6 | N447/24 | Kv11.3 K+ channel |
| L61C/30 | Kv2.1 K+ channel | N71/37 | HCN2 | N271/44 | ASIC1 | N448/88 | Kv8.2 K+ channel |
| L62/29 | Copper ATPase 2 (Wilson's disease protein) | N72/16 | Kv3.4 K+ channel | N274/8 | Synaptotagmin-5 | N449/73 | VMAT2 |
| L64/32 | Kv1.2 K+ channel | N74/25 | TrpM7 | N275/14 | Synaptotagmin-7 | N450/53* | BDNF |
| L71/5 | Kv1.4 K+ channel (extracellular) | N75/3 | mGluR1/5 (Group I) glutamate receptor | N276A/15 | Synaptotagmin-9 | N451/73 | ICK |
| L71/22 | Kv1.4 K+ channel (extracellular) | N75/33 | mGluR1/5 (Group I) glutamate receptor | N283/7 | Lgi1 | N452/30 | GABA(A)R, Gamma2L/S |
| L76/36 | Kv1.2 K+ channel | N77/15 | TrpC4 | N286/74 | Zebrafish PSD Marker | N452/69 | GABA(A)R, Gamma2L |

| | | | | | | | |
|---|---|---|---|---|---|---|---|
| L80/21 | Kv2.1 K+ channel | N81/2 | GABA(B)R2 | N290B/25 | ARHGAP4 | N452/73 | GABA(A)R, Gamma2L/S |
| L83/11 | Kv2.1 K+ channel | N86/20 | GFP | N294A/10 | Brevican | N452/81 | GABA(A)R, Gamma2L/S |
| L83/81 | Kv2.1 K+ channel | N86/44 | GFP | N294A/6 | Brevican | N454/91 | ANO3/TMEM16C |
| L86/2 | AMIGO-1 | N86/8 | GFP | N295B/54 | Arl13b | N455/15 | Kir3.3 K+ channel |
| L86/14 | AMIGO-1 | N87/25 | GABA(A)R, Beta3 | N302/10 | BBS3/Arl6 | N456/39 | Proser1 |
| L86/33* | AMIGO-1 | N92/14 | FGF11/FHF3 | N304B/115 | BBS5 | N458/10 | Kv6.4 K+ channel |
| L86/36 | AMIGO-1 | N95/35 | GABA(A)R, Alpha1 | N307/12 | Histone H3-acetyl-Lys56 | N459/84 | SAPAP2 |
| L100/1 | Kv2.1 K+ channel pS586 | N98/47 | Neuroligin-4 | N308/48 | GluN1/NR1 glutamate receptor | N460/19 | Kv9.2 K+ channel |
| L102/45 | SynDIG4/Prrt1 | N100/13 | GST | N309A/21 | Histone H4-dimethyl-Arg3 | N461/19 | Kv9.1 K+ channel |
| L106/4 | Gephyrin | N104/37 | SNAT1 | N312/10 | NSD1 | N463/52* | DEPDC5 |
| L106/22 | Gephyrin | N105/13 | Ankyrin-B | N319A/14 | SUR2A | N465/11 | Kir2.4 K+ channel |
| L106/23 | Gephyrin | N105/17 | Ankyrin-B | N323A/31 | SUR1 and SUR2B | N467/1 | TRESK potassium channel |
| L106/83 | Gephyrin | N106/20 | Ankyrin-G | N324/2 | Rem2 | N468/37 | THIK-2 potassium channel |
| L107/39 | Neuroligin-2 | N106/36* | Ankyrin-G | N325B/65 | THAP1 | N470/22 | TRPM6 |
| L107/90* | Neuroligin-2 | N106/65 | Ankyrin-G (staining) | N326D/2 | REEP2 | N471/27 | Kv1.7 K+ channel |
| L107/93 | Neuroligin-2 | N112/16 | Kir2.1 K+ channel | N326D/13 | REEP | N472/88 | MiRP4 K+ channel |
| L107/95 | Neuroligin-2 | N112B/14 | Kir2.1 K+ channel | N327/95 | GluN2A/NR2A glutamate receptor | N473/36 | Cavbeta3 Ca2+ channel |
| L109/39 | Calbindin | N112B/29 | Kir2.1 K+ channel | N327A/38 | GluN2A/NR2A glutamate receptor | N476/9 | ZIP3 |
| L109/57 | Calbindin | N119/44 | MPP8 | N330A/80 | Alpha-2C adrenergic receptor | N479/107 | VAPA/B |
| L109/62 | Calbindin | N121A/1 | Pannexin-2 | N332B/15 | BAF53b | N479/12 | VAPA |
| L113/13 | Homer1L | N123/19 | Histone H3-pThr11 | N336B/83 | BAF53a | N479/22 | VAPA |
| L113/26 | Homer1L | N123/48 | Histone H3-pThr11 | N341/12 | LRRK1 | N483/26 | Rufy3 |
| L113/27 | Homer1L | N125/10 | Thorase/Atad1 | N343/26 | NrCAM | N483/84 | Rufy3 |
| L113/28 | Homer1L | N126B/31 | Neuregulin-CRD (Cysteine-rich domain, Type III) | N345/51 | REEP1 | N483/126 | Rufy3 |
| L113/29 | Homer1L | N128A/2 | Haspin/GSG2 | N347/42 | ATAT1 | N485/22 | Prrt2 |
| L113/44 | Homer1L | N128A/4 | Maltose binding protein (MBP) | N349/96 | Foxi2 | N486/25 | c-Fos |
| L113/62 | Homer1L/S | N129/15 | Mad3 (human) | N356/9 | SVOP | N486/32 | c-Fos |
| L113/130 | Homer1L/S | N132A/12 | KLH | N357/6* | PINK1 | N486/76 | c-Fos |

**Supplementary Table 2.** List of R-mAbs and their targets that were successfully cloned into the IgG2a mammalian expression plasmid using the sequence-based Gibson Assembly cloning method. * cloned from hybridomas with more than two $V_L$ sequences. ** cloned from a hybridoma with two $V_L$ and two $V_H$ sequences.
