## Supplementary Table 3 for "NeuroMabSeq: high volume acquisition, processing, and curation of hybridoma sequences and their use in generating recombinant monoclonal antibodies and scFvs for neuroscience research"

### Supplementary Table 3. List of R-mAbs successfully transferred to IgG subclass-switched expression plasmids.

| Transferred to IgG2b Expression Plasmid | | | | Transferred to IgG1 Expression Plasmid | |
|---|---|---|---|---|---|
| R-mAb | Target | R-mAb | Target | R-mAb | Target |
| A12/18 | Pan-Neurofascin (extracellular) | N59/20 | GluN2B/NR2B glutamate receptor | A12/18 | Pan-Neurofascin (extracellular) |
| K7/45 | Kv1.5 K+ channel | N59/36 | GluN2B/NR2B glutamate receptor | K20/78 | Kv1.1 K+ channel |
| K13/31 | Kv1.4 K+ channel | N68/6 | Nav1.7 Na+ channel | K28/43 | PSD-95 |
| K14/16 | Kv1.2 K+ channel | N69/46 | Shank3 | K57/1 | Kv4.2 K+ channel (external) |
| K17/70 | Kvbeta2 K+ channel | N70/28 | HCN1 | K65/35 | CASPR/Neurexin IV |
| K20/78 | Kv1.1 K+ channel | N76/8 | Ataxin-1, 11NQ | K75/41 | Kv4.3 K+ channel |
| K28/43 | PSD-95 | N86/38 | GFP | L108/92 | VIP |
| K39/25 | Kv2.1 K+ channel (external) | N86/8 | GFP | L109/39 | Calbindin |
| K57/1 | Kv4.2 K+ channel (external) | N97A/31 | Neuroligin-1 | L113/130 | Homer1L/S |
| K64/15 | SAP97 | N106/36 | Ankyrin-G | L113/27 | Homer1L |
| K65/35 | CASPR/Neurexin IV | N110/29 | Neuroligin-3 | L118/80 | VGAT |
| K66/38 | KChIP3 K+ channel | N112B/14 | Kir2.1 K+ channel | L122/6 | Calretinin |
| K74/71 | Nav1.1 Na+ channel | N114/10 | HCN4 | L122/68 | Calretinin |
| K75/41 | Kv4.3 K+ channel | N133/21 | RGS14 | L127/12 | GAD65 + GAD67 |
| K89/34 | Kv2.1 K+ channel | N133/35 | RGS14 | L21/32 | GluA2/GluR2 glutamate receptor |
| L6/60 | Slo1/BKAlpha maxi-K+ channel | N134/12 | Nav1.8 Na+ channel | L6/60 | Slo1/BKAlpha maxi-K+ channel |
| L21/32 | GluA2/GluR2 glutamate receptor | N149/25 | Olig1 | L86/33 | AMIGO-1 |
| L28/4 | Kv4.2 K+ channel | N151/3 | GABA(A)R, Delta | N1/12 | KCC2 |
| L57/46 | Cav1.2/1.3 Ca2+ channel | N168/6 | Navbeta4 Na+ channel | N86/8 | Nav1.7 Na+ channel |
| L60/4 | Copper ATPase 1 (Menke's disease protein) | N170A/1 | Neurexin-1-Beta | N106/36 | Ankyrin-G |
| L86/33 | AMIGO-1 | N178A/9 | Cav3.1 Ca2+ channel | N112B/14 | Kir2.1 K+ channel |
| L86/36 | AMIGO-1 | N182/17 | QKI-6 | N133/21 | RGS14 |
| L106/83 | Gephyrin | N196/16 | PARIS/ZNF746 | N134/12 | Nav1.8 Na+ channel |
| L107/39 | Neuroligin-2 | N201/35 | Iduna/RNF146 | N149/25 | Olig1 |
| L108/92 | VIP | N206A/8 | GFAP | N170A/1 | Neurexin-1-Beta |
| L109/39 | Calbindin | N212/17 | TRIP8b (constant) | N206A/8 | GFAP |
| L113/27 | Homer1L | N241A/34 | LRRK2/Dardarin, C-terminus | N241A/34 | LRRK2/Dardarin, C-terminus |
| L113/130 | Homer1L/S | N244/5 | SynCAM4 | N263/31 | Cav1.2 Ca2+ channel |
| L114/3 | Parvalbumin | N263/31 | Cav1.2 Ca2+ channel | N28/9 | VGluT1 |
| L115/13 | NPY/Neuropeptide Y | N289/16 | SUR1 | N29/29 | VGluT2 |
| L118/80 | VGAT | N295B/66 | Arl13b | N295B/66 | Arl13b |
| L122/6 | Calretinin | N308/48 | GluN1/NR1 glutamate receptor | N308/48 | GluN1/NR1 glutamate receptor |
| L122/68 | Calretinin | N327/95 | GluN2A/NR2A glutamate receptor | N327/95 | GluN2A/NR2A glutamate receptor |
| L127/8 | GAD67 | N355/1 | GluA1/GluR1 glutamate receptor | N355/1 | GluA1/GluR1 glutamate receptor |
| L127/12 | GAD65/67 | N410/17 | Kv3.2 K+ channel | N410/17 | Kv3.2 K+ channel |
| N1/12 | KCC2 | N425/45 | VAChT | N425/45 | VAChT |
| N3/26 | KCNT1/Slo2.2/Slack K+ channel | | | N52A/42 | Mortalin/GRP75 |
| N6/38 | VAChT | | | N59/20 | GluN2B/NR2B glutamate receptor |
| N7/18 | Cavbeta1 Ca2+ channel | | | N59/36 | GluN2B/NR2B glutamate receptor |
| N11/33 | KCNT2/Slo2.1/Slick K+ channel | | | N6/38 | VAChT |
| N28/9 | VGlut1 | | | N68/6 | Nav1.7 Na+ channel |
| N29/29 | VGluT2 | | | N70/28 | HCN1 |
| N52A/42 | Mortalin/GRP75 | | | N86/38 | GFP |

**Supplementary Table 3.** List of R-mAbs and their targets that were successfully transferred from the IgG2a mammalian expression plasmid into either an IgG2b and/or IgG1 mammalian expression plasmid using restriction digest/ligation-based cloning.
