## Supplementary Table 4 for "NeuroMabSeq: high volume acquisition, processing, and curation of hybridoma sequences and their use in generating recombinant monoclonal antibodies and scFvs for neuroscience research"

### Supplementary Table 4. List of scFvs successfully cloned via sequence-based cloning.

| scFv | Target | Plasmid | Tags | scFv | Target | Plasmid | Tags |
|---|---|---|---|---|---|---|---|
| 1D8 | LRRK2/Dardarin-pSer910 | pcDNA3.4 | HA<br>sortase<br>His | N67/15 | TrpC5 | pcDNA3.4 | HA<br>sortase<br>His |
| A12/18 | Pan-Neurofascin (extracellular) | pcDNA3.1 | sortase<br>His | N68/6 | Nav1.7 Na channel | pcDNA3.1 | sortase<br>His |
| A12/18 | Pan-Neurofascin (extracellular) | pcDNA3.1 | HA<br>sortase<br>His | N69/46 | Shank3 | pcDNA3.4 | HA<br>sortase<br>His |
| K7/45 | Kv1.5 K channel | pcDNA3.4 | HA<br>sortase<br>His | N86/20 | GFP | pcDNA3.4 | HA<br>sortase<br>His |
| K14/16 | Kv1.2 K channel | pcDNA3.1 | HA<br>His | N86/38 | GFP | pcDNA3.1 | HA<br>His |
| K17/70 | Kvbeta2/KCNAB2 K channel | pcDNA3.4 | HA<br>sortase<br>His | N86/44 | GFP | pcDNA3.4 | HA<br>sortase<br>His |
| K20/78 | Kv1.1 K channel | pcDNA3.1 | sortase<br>His | N93A/49 | GABA(B)R1 | pcDNA3.4 | HA<br>sortase<br>His |
| K20/78 | Kv1.1 K channel | pcDNA3.1 | HA<br>sortase<br>His | N100/13 | GST | pcDNA3.4 | HA<br>sortase<br>His |
| K28/38 | PSD-95 | pcDNA3.4 | HA<br>sortase<br>His | N103/31 | Aldh1L1 | pcDNA3.1 | sortase<br>His |
| K28/42 | PSD-95 PSD-93 | pcDNA3.4 | HA<br>sortase<br>His | N103/31 | Aldh1L1 | pcDNA3.1 | HA<br>sortase<br>His |
| K28/43 | PSD-95 | pCDNA3.4 | HA<br>His | N105/13 | Ankyrin-B | pcDNA3.4 | HA<br>sortase<br>His |
| K28/43 | PSD-95 | pcDNA3.1 | HA<br>Flag<br>sortase | N105/17 | Ankyrin-B | pcDNA3.4 | HA<br>sortase<br>His |
| K28/74 | PSD-95 | pcDNA3.4 | HA<br>sortase<br>His | N106/20 | Ankyrin-G | pcDNA3.4 | HA<br>sortase<br>His |
| K39/25 | Kv1.2 K channel (extracellular) | pCDNA3.4 | HA<br>His | N114/10 | HCN4 | pcDNA3.1 | HA<br>sortase<br>His |
| K56A/50 | CASK | pcDNA3.4 | HA<br>sortase<br>His | N125/10 | Thorase/Atad1 | pcDNA3.4 | HA<br>sortase<br>His |
| K56A/57 | CASK | pcDNA3.4 | HA<br>sortase<br>His | N126B/31 | Neuregulin-CRD (Cysteine-rich domain, Type III) | pcDNA3.4 | HA<br>sortase<br>His |
| K60/73 | KChIP2b K channel | pcDNA3.4 | HA<br>sortase<br>His | N129/15 | Mad3 (human) | pcDNA3.4 | HA<br>sortase<br>His |
| K65/35 | CASPR/Neurexin IV | pcDNA3.1 | sortase<br>His | N133/21 | RGS14 | pcDNA3.1 | sortase<br>His |
| K66/27 | KChIP3 K channel | pcDNA3.4 | HA<br>sortase<br>His | N133/35 | RGS14 | pcDNA3.4 | HA<br>sortase<br>His |
| K66/38 | KChIP3 K channel | pcDNA3.4 | HA<br>sortase<br>His | N134/12 | Nav1.8 Na channel | pcDNA3.1 | sortase<br>His |
| K66/59 | KChIP3 K channel | pcDNA3.4 | HA<br>sortase<br>His | N134/12 | Nav1.8 Na channel | pcDNA3.1 | HA<br>sortase<br>His |
| K89/34 | Kv2.1 K channel | pCDNA3.4 | HA<br>His | N144/17 | Gamma-protocadherin-A3 | pcDNA3.4 | HA<br>sortase<br>His |

| | | | | | | | |
|---|---|---|---|---|---|---|---|
| K89/34 | Kv2.1 K channel | pcDNA3.1 | HA<br>Flag<br>sortase | N144/32 | Pan-Gamma-protocadherin-A | pcDNA3.4 | HA<br>sortase<br>His |
| K89/34<br>Flag | Kv2.1 K channel | pcDNA3.4 | HA<br>His<br>Flag | N149/25 | Olig1 | pcDNA3.4 | HA<br>sortase<br>His |
| K89/41 | Kv2.1 K channel | pcDNA3.4 | HA<br>sortase<br>His | N151/3 | GABA(A)R, Delta | pcDNA3.4 | HA<br>sortase<br>His |
| K96/7 | Pancortin | pcDNA3.4 | HA<br>sortase<br>His | N159/5 | Pan-Gamma-protocadherin-Constant | pcDNA3.4 | HA<br>sortase<br>His |
| L11A/41 | Pan-Neurofascin | pcDNA3.4 | HA<br>sortase<br>His | N160/21 | SynDIG3/Tmem91 | pcDNA3.4 | HA<br>sortase<br>His |
| L23/27 | Kv1.3 K channel | pcDNA3.1 | HA<br>sortase<br>His | N170A/1 | Neurexin-1-Beta | pcDNA3.1 | HA<br>sortase<br>His |
| L24/1 | IP3 receptor, type 1 | pcDNA3.4 | HA<br>sortase<br>His | N170A/26 | Neurexin-1-Beta | pcDNA3.4 | HA<br>sortase<br>His |
| L48A/9 | Cav1.3 Ca2 channel | pcDNA3.4 | HA<br>sortase<br>His | N183/15 | QKI-7 | pcDNA3.4 | HA<br>sortase<br>His |
| L61C/30 | Kv2.1 K channel | pcDNA3.4 | HA<br>sortase<br>His | N185/7 | NALCN | pcDNA3.4 | HA<br>sortase<br>His |
| L71/5 | Kv1.4 K channel (extracellular) | pcDNA3.4 | HA<br>sortase<br>His | N196/16 | PARIS/ZNF746 | pcDNA3.4 | HA<br>sortase<br>His |
| L76/36 | Kv1.2 K channel | pcDNA3.4 | HA<br>sortase<br>His | N201/35 | Iduna/RNF146 | pcDNA3.4 | HA<br>sortase<br>His |
| L83/11 | Kv2.1 K channel | pcDNA3.4 | HA<br>sortase<br>His | N206A/8 | GFAP | pcDNA3.1 | sortase<br>His |
| L86/33 | AMIGO-1 | pcDNA3.4 | HA<br>sortase<br>His | N207/27 | LRP4 (extracellular) | pcDNA3.4 | HA<br>sortase<br>His |
| L107/39 | Neuroligin-2 | pcDNA3.1 | sortase<br>His | N212/17 | TRIP8b (constant) | pcDNA3.4 | HA<br>sortase<br>His |
| L107/90 | Neuroligin-2 | pcDNA3.4 | HA<br>sortase<br>His | N221/17 | TRIP8b (constant) | pcDNA3.1 | sortase<br>His |
| L107/95 | Neuroligin-2 | pcDNA3.4 | HA<br>sortase<br>His | N212A/34 | TRIP8b (exon 1b) | pcDNA3.4 | HA<br>sortase<br>His |
| L109/57 | Calbindin | pCDNA3.4 | HA<br>His | N221/17 | TrpV1 | pcDNA3.4 | HA<br>sortase<br>His |
| L113/13 | Homer1L | pcDNA3.1 | sortase<br>His | N225A/10 | FGF13/FHF2, B isoform | pcDNA3.4 | HA<br>sortase<br>His |
| L113/13 | Homer1L | pcDNA3.1 | HA<br>sortase<br>His | N241A/34 | LRRK2/Dardarin, C-terminus | pcDNA3.1 | sortase<br>His |
| L113/130 | Homer1L/S | pcDNA3.1 | sortase<br>His | N263/31<br>LC1 | Cav1.2 Ca2 channel | pCDNA3.4 | HA<br>His |
| L113/130 | Homer1L/S | pcDNA3.1 | HA<br>sortase<br>His | N270/47 | Synaptotagmin-6 | pcDNA3.4 | HA<br>sortase<br>His |
| L113/27 | Homer1L | pcDNA3.1 | sortase<br>His | N276A/15 | Synaptotagmin-9 | pcDNA3.4 | HA<br>sortase<br>His |
| L113/27 | Homer1L | pcDNA3.1 | HA<br>sortase<br>His | N289/16 | SUR1 | pcDNA3.4 | HA<br>sortase<br>His |

| | | | | | | | |
|---|---|---|---|---|---|---|---|
| L113/28 | Homer1L | pcDNA3.4 | HA sortase His | N295B/66 | Arl13b | pcDNA3.1 | HA His |
| L113/44 | Homer1L | pcDNA3.4 | HA sortase His | N355/1 | GluA1/GluR1 glutamate receptor | pcDNA3.1 | sortase His |
| L114/3 | Parvalbumin | pCDNA3.4 | HA His | N363/71 | Kir6.2 K channel | pcDNA3.4 | HA sortase His |
| L114/3 Flag | Parvalbumin | pcDNA3.4 | HA His Flag | N366/60 | Kir6.1 K channel | pcDNA3.4 | HA sortase His |
| L114/81 | Parvalbumin | pCDNA3.4 | HA His | N367/51 | Shank1 and Shank3 | pcDNA3.4 | HA sortase His |
| L115/13 | NPY/Neuropeptide Y | pcDNA3.1 | sortase His | N372B/1 | Kv2.2 K channel | pcDNA3.4 | HA sortase His |
| L115/13 | NPY/Neuropeptide Y | pcDNA3.1 | HA sortase His | N372B/60 | Kv2.2 K channel | pcDNA3.4 | HA sortase His |
| L117/1 | IRSp53/BAIAP2 | pcDNA3.1 | sortase His | N375/67 | Kv3.3 K channel | pcDNA3.4 | HA sortase His |
| L117/1 | IRSp53/BAIAP2 | pcDNA3.1 | HA sortase His | N393/2 | Beta4-spectrin | pcDNA3.4 | HA sortase His |
| L118/14 | VGAT | pcDNA3.4 | HA sortase His | N398A/34 | GABA(A)R, Alpha4 | pcDNA3.4 | HA sortase His |
| L120/12 | Collybistin | pcDNA3.1 | sortase His | N399/19 | GABA(A)R, Alpha2 | pcDNA3.4 | HA sortase His |
| L121/133 | nNOS/NOS1 | pcDNA3.4 | HA sortase His | N402/46 | Thyroid hormone receptor beta1 | pcDNA3.4 | HA sortase His |
| L121/136 | nNOS/NOS1 | pcDNA3.4 | HA sortase His | N403/63 | Pan-Thyroid hormone receptor | pcDNA3.4 | HA sortase His |
| L122/6 | Calretinin | pCDNA3.4 | HA His | N414/25 | NCKX4 | pcDNA3.4 | HA sortase His |
| L122/68 | Calretinin | pcDNA3.1 | sortase His | N422/18 | GluN2C/NR2C glutamate receptor | pcDNA3.4 | HA sortase His |
| L124/59 | Bassoon | pcDNA3.1 | sortase His | N432/21 | VSP | pcDNA3.4 | HA sortase His |
| L124/59 | Bassoon | pcDNA3.1 | HA sortase His | N432/63 | VSP | pcDNA3.4 | HA sortase His |
| L125/129 | Pan-Synapsin | pcDNA3.1 | sortase His | N448/88 | Kv8.2 K channel | pcDNA3.4 | HA sortase His |
| L127/8 | GAD67 | pcDNA3.1 | sortase His | N449/73 | VMAT2 | pcDNA3.4 | HA sortase His |
| L130/1 | Tiam1 | pcDNA3.4 | HA sortase His | N452/73 | GABA(A)R, Gamma2L/S | pcDNA3.1 | sortase His |
| L131/17 | SPHKAP | pcDNA3.4 | HA sortase His | N452/73 | GABA(A)R, Gamma2L/S | pcDNA3.1 | HA sortase His |
| L131/20 | SPHKAP | pcDNA3.4 | HA sortase His | N452/81 | GABA(A)R, Gamma2L/S | pcDNA3.1 | sortase His |
| L131/27 | SPHKAP | pcDNA3.4 | HA sortase His | N452/81 | GABA(A)R, Gamma2L/S | pcDNA3.1 | HA sortase His |

| | | | | | | | |
|---|---|---|---|---|---|---|---|
| N1/12 | KCC2 | pcDNA3.1 | HA sortase His | N471/27 | Kv1.7 K channel | pcDNA3.4 | HA sortase His |
| N16B/8 | Kv3.1b K channel | pcDNA3.1 | sortase His | N479/107 | VAPA/B | pcDNA3.4 | HA sortase His |
| N18/28 | PSD-93/Chapsyn-110 | pcDNA3.4 | HA sortase His | N479/12 | VAPA | pcDNA3.4 | HA sortase His |
| N22/21 | Shank1 | pcDNA3.4 | HA sortase His | N483/26 | Rufy3 | pcDNA3.4 | HA sortase His |
| N23B/6 | Shank2 | pcDNA3.4 | HA sortase His | N483/84 | Rufy3 | pcDNA3.4 | HA sortase His |
| N23B/49 | Pan-Shank | pcDNA3.1 | HA sortase His | N483/126 | Rufy3 | pcDNA3.1 | sortase His |
| N28/9 | VGlut1 | pcDNA3.1 | HA His | N483/126 | Rufy3 | pcDNA3.1 | HA sortase His |
| N28/9 Flag | VGlut1 | pcDNA3.4 | HA His Flag | N486/32 | c-Fos | pCDNA3.4 | HA His |
| N29/29 | VGlut2 | pcDNA3.1 | HA His | N486/76 | c-Fos | pcDNA3.4 | HA sortase His |
| N43/6 | Kv7.4/KCNQ4 K channel | pcDNA3.4 | HA sortase His | N112B/29 | Kir2.1 K+ channel | pcDNA3.4 | HA sortase His |
| N53/32 | BKbeta2 K channel | pcDNA3.4 | HA sortase His | N140A/12 | SALM5/LRFN5 | pcDNA3.4 | HA sortase His |
| N52A/42 | Mortalin/GRP75 | pcDNA3.1 | HA His | N155/9 | Uncx | pcDNA3.4 | HA sortase His |
| N52B/27 | SALM2/LRFN1 | pcDNA3.4 | HA sortase His | N294A/6 | Brevican | pcDNA3.4 | HA sortase His |
| N59/20 | GluN2B/NR2B glutamate receptor | pcDNA3.1 | sortase His | N312/10 | NSD1 | pcDNA3.4 | HA sortase His |
| N59/20 | GluN2B/NR2B glutamate receptor | pcDNA3.1 | HA sortase His | N411/51 | Arx | pcDNA3.4 | HA sortase His |
| N59/36 | GluN2B/NR2B glutamate receptor | pcDNA3.1 | HA sortase His | N416/57 | GluN3A/NR3A glutamate receptor | pcDNA3.4 | HA sortase His |
| N59/36 | GluN2B/NR2B glutamate receptor | pcDNA3.1 | HA His | N424/45 | Glycine receptor Alpha3 | pcDNA3.4 | HA sortase His |
| D4/154 | Kv2.1 K+ channel | pcDNA3.4 | HA sortase His | N428/12 | WRN | pcDNA3.4 | HA sortase His |
| K57/41 | Kv4.2 K+ channel (external) | pcDNA3.4 | HA sortase His | N452/30 | GABA-A-R-Gamma2L/S, GABRG2 | pcDNA3.4 | HA sortase His |
| L24/21 | IP3R | pcDNA3.4 | HA sortase His | N473/36.3 | Cavbeta3 Ca2+ channel | pcDNA3.4 | HA sortase His |
| L86/14 | AMIGO-1 | pcDNA3.4 | HA sortase His | N485/22 | Prrt2 | pcDNA3.4 | HA sortase His |
| L86/2 | AMIGO-1 | pcDNA3.4 | HA sortase His | **$V_L$-linker-$V_H$** | | | |
| L106/23 | Gephyrin | pcDNA3.4 | HA sortase His | L6/60 | Slo1/BKAlpha maxi-K channel | pcDNA3.4 | HA sortase His |

| | | | | | | | |
|---|---|---|---|---|---|---|---|
| L114/38 | Parvalbumin | pcDNA3.4 | HA sortase His | N86/8 | GFP | pcDNA3.4 | HA sortase His |
| L122/144 | Calretinin | pcDNA3.4 | HA sortase His | N95/35 | GABA(A)R, Alpha1 | pcDNA3.4 | HA sortase His |
| L123/46 | GABA(A)R, Theta | pcDNA3.4 | HA sortase His | N106/36 | Ankyrin-G | pcDNA3.4 | HA sortase His |
| N7/18 | Cavbeta1 Ca2+ channel | pcDNA3.4 | HA sortase His | N112B/14 | Kir2.1 K channel | pcDNA3.4 | HA sortase His |
| N56/21 | FGF14a | pcDNA3.4 | HA sortase His | N377/20 | MAP3K12 | pcDNA3.1 | HA sortase His |
| N112/16 | Kir2.1 K+ channel | pcDNA3.4 | HA sortase His | | | | |

**Supplementary Table 4.** List of scFvs and their targets that were successfully cloned into either pcDNA3.1 or pcDNA3.4 mammalian expression plasmids using the sequence-based Gibson Assembly cloning method. All are in the $V_H$-linker-$V_L$ format except as noted. Tags indicate the presence of HA or Flag epitope tags, a sortase labelling tag and/or the 6xHis purification tag at the C-terminus of the scFv.
